# At what spatial scales are alternative stable states relevant in highly interconnected ecosystems?

**DOI:** 10.1101/717504

**Authors:** Vadim A. Karatayev, Marissa L. Baskett

**Affiliations:** Department of Environmental Science and Policy, University of California, Davis; Graduate Group in Ecology, University of California, Davis

**Keywords:** Alternative stable states, ecological resilience, dispersal, disturbance, kelp forest, urchin barren, stochastic population dynamics, temperate rocky reefs

## Abstract

Whether ecosystems recover from disturbance depends on the presence of alternative stable states, which are theoretically possible in simple models of many systems. However, definitive empirical evidence for this phenomenon remains limited to demographically closed ecosystems such as lakes. In more interconnected systems such as temperate rocky reefs, the local relevance of alternative stable states might erode as immigration overwhelms local feedbacks and produces a single stable state. At larger spatial scales, dispersal might counter localized disturbance and feedbacks to synchronize states throughout a region. Here, we quantify how interconnectedness affects the relevance of alternative stable states using dynamical models of California rocky reef communities that incorporate observed environmental stochasticity and feedback loops in kelp-urchin-predator interactions. Our models demonstrate the potential for localized alternative states despite high interconnectedness likely due to feedbacks affecting dispersers as they settle into local communities. Regionally, such feedbacks affecting settlement can produce a mosaic of alternative stable states that span local (10-20km) scales despite the synchronizing effect of long-distance dispersal. The specific spatial scale and duration of each state predominantly depend on the scales of environmental variation and on local dynamics (here, fishing). Model predictions reflect observed scales of community states in California rocky reefs and suggest how alternative states co-occur in the wide array of marine and terrestrial systems with settlement feedbacks.

## Introduction

When qualitatively distinct, alternatively stable ecosystem states occur under the same environmental conditions, brief (“pulse”) disturbances can lead to abrupt and persistent ecological shifts (Scheffer et al. 2001). In such cases, the persistence of a given ecological state depends on the magnitude of perturbations that ecosystems can withstand and return to the target state (‘ecological resilience’ hereafter; Holling 1973). Models and empirically-observed feedbacks reinforcing distinct states suggest that alternative stable states could, theoretically, underlie observed shifts following environmental change in many ecosystems, including the collapse of fish stocks, declines in foundational species including corals and kelp, replacement of forests by fire-prone grasslands, and algal dominance in lakes (Scheffer et al. 2009). Alternatively, such state shifts may be readily reversed if only one state is stable under a given set of environmental conditions, where that state can still rapidly change with the environment. Therefore, in addition to informing a basic understanding of the structure of ecological communities, resolving whether or not state shifts represent alternative stable states can affect conservation management decisions such as the degree of restoration necessary for recovery of a target state (Suding et al. 2004). However, empirically establishing the relevance of alternative stable states requires long-term data of each state at large spatial scales under identical environmental conditions and therefore remains debated in most systems (Petraitis and Dudgeon 2004; Petraitis 2013).

One of the factors that might affect the relevancy of alternative stable states to many systems is demographic interconnectedness (Petraitis and Latham 1999; Petraitis and Dudgeon 2004). In particular, inputs of individuals from outside areas may overwhelm the feedbacks maintaining alternative stable states at small spatial scales and induce a “rescue effect” (Brown and Kodric-Brown 1977) that always allows eventual recovery of the original state after disturbance. Consequently, the best evidence for the existence of alternative stable states and their relevance to management remains limited to relatively small, demographically closed shallow lake ecosystems (Scheffer et al. 1993; Schroder et al. 2005). By contrast, in most terrestrial and marine systems manipulative studies occur on small scales (e.g., *<*100 m, McGill 2010) while many animal taxa disperse above the scale of kilometers (Kinlan and Gaines 2003), which can dynamically link communities across tens to hundreds of kilometers (Gouhier et al. 2010).

At the larger spatial scale of an entire ecosystem, high interconnectedness among locations raises the question of whether alternative stable states, if relevant, manifest within local communities or at system-wide scales. On the one hand, localized disturbances induce ecosystem heterogeneity that can maintain localized alternative stable states: for instance, in models with Allee effects, demographic stochasticity in a given year can reinforce population persistence in some habitat patches and promote extinction in others (Martin et al. 2015). Analogous heterogeneity in savannas, coral reefs, and predator-prey communities might maintain localized alternative stable states in spite of high external input (Durrett and Levin 1994; Mumby et al. 2007; Schertzer et al. 2015). On the other hand dispersal at the ecosystem level can synchronize ecological states across space and produce alternative stable states at system-wide scales. In particular, spatial models emphasize that under high levels of dispersal sudden collapses to undesired ecosystem states can propagate across large scales in a domino effect (van Nes and Scheffer 2005; Hughes et al. 2005, 2013), with recovery possible only through large-scale restoration efforts (van de Leemput et al. 2015). Resolving the expected scale of alternative stable states and its drivers informs the scales at which empirical efforts could detect this phenomenon, the socioeconomic consequences of ecosystem collapses to undesired states, and which management approaches might effectively maintain desired states.

Temperate rocky reef ecosystems throughout the world exemplify systems with both extensive dispersal and a hypothesized potential for alternative stable states. Two observed ecological states of these communities are kelp forests with abundant urchin predators and barrens where urchins overgraze kelp and predators are rare (reviewed in Lawrence 1975; Steneck et al. 2002; Konar and Estes 2003; Ling et al. 2015). When fishing reduces predator densities, transitions between these states often follow localized, stochastic events such as pulses of urchin larvae and kelp loss during intense storms (Hart and Scheibling 1988; Cavanaugh et al. 2011). These community shifts typically occur on small scales (≤ 20 km of coastline) and can persist for years to decades (reviewed in Filbee-Dexter and Scheibling 2014). While a number of feedbacks have the potential to drive alternative stable states on temperate reefs, such as when abundant kelp facilitate recruitment of urchin predators (Ling et al. 2015) and barrens facilitate urchin recruitment (Baskett and Salomon 2010), urchin barrens might instead represent gradual recoveries from disturbances given high dispersal and connectivity. Specifically, urchin and predator dispersal on 50-100 km scales (Waples and Rosenblatt 1987; Edmands et al. 1996) largely decouples larval supply from local consumer densities (Okamoto 2014), and might rescue predators from local disturbance (Connell and Sousa 1983) or propagate and synchronize urchin barrens over large scales (Hughes et al. 2005). Where shifts to urchin barrens have occurred, resolving the relevance and scale of alternative stable states can inform whether reduction in predator harvest alone allows kelp forest recovery, or whether additional restoration efforts (e.g., urchin harvest; House et al. 2017) might be necessary at specific scales.

Here, we quantify whether and at what spatial scales alternative stable states and the associated concept of ecological resilience can be relevant in temperate rocky reefs given long-distance dispersal. We develop a tritrophic community model with empirically grounded feedback mechanisms that can drive alternative forested and barren states by regulating settlement success. We first use a 1-patch model to explore how the fraction of external larval supply affects the presence of alternative stable states, and then resolve their spatial and temporal scales using a spatially explicit model formulation that incorporates observed stochasticity. Finally, we compare the predicted scales of the alternative states from our model to long-term data. From these analyses, we find a potential for local relevance of alternative stable states in systems with high connectivity and quantify how feedbacks and local disturbance interact to determine community structure across locations and through time.

## Materials and methods

### Study system

We base our analysis on temperate rocky reefs in the northern California Channel Islands, which represent a c.a. 300 km total coastline with well-documented shifts between distinct community states (Filbee-Dexter and Scheibling 2014). Like many temperate rocky reefs around the world, strong trophic interactions characterize this system, and primarily involve kelp (*Macrocystis pyrifera*), sea urchin herbivores (*Strongylocentrotus purpuratus*), and urchin predators: sheephead (*Semicossyphus pulcher*) and spiny lobsters (*Panulirus interruptus*) (Filbee-Dexter and Scheibling 2014; Hamilton and Caselle 2015). Kelp facilitate urchin predators by providing shelter that can double survival rates of predator larvae compared to exposed areas (Smith and Herrkind 1992; Lipcius et al. 1998, Appendix S1: Section S1). Predator recruitment facilitation by kelp creates a strong feedback often invoked by empirical studies of temperate rocky reefs (Ling et al. 2015) that can produce alternatively stable forest and barren states under intensive predator fishing in closed-system models (Marzloff et al. 2013; Appendix S1: Section S2, Fig. S5a). Note that predator recruitment facilitation, the focus here, is one of many possible feedbacks hypothesized to drive alternative stable states in kelp systems (Ling et al. 2015); additional possible mechanisms include recruitment facilitation of urchins to crustose coralline algae that increases in barrens (Baskett and Salomon 2010) and intensified urchin foraging behavior when kelp (Ebeling et al. 1985; Harrold and Reed 1985) or predators (Cowen 1983) are rare.

Local communities in this system are highly interconnected because dispersal distances greatly exceed the scale of predominant environmental stochasticity. Although adult consumer home ranges and kelp dispersal occur within rocky kelp habitat patches (1-5 km of coastline; Topping et al. 2005; Castorani et al. 2015), consumer larvae disperse over much longer distances (50-100 km on average; Table 1). We consider two sources of local-scale stochasticity that regulate community composition and can form persistent urchin barrens: winter storms, which annually lead to a 70% loss of kelp biomass (Dayton et al. 1999; Cavanaugh et al. 2011), and urchin larval survival, which varies temporally by an order of magnitude (Hart and Scheibling 1988; Schroeter et al. 1996). As with our focal feedback mechanism, these are not the only possible sources of environmental stochasticity, which also include extreme warm water events leading to kelp loss (Dayton et al. 1999) and disease outbreaks affecting predators (Harvell et al. 2019) and urchins (Ling et al. 2015).

**Table 1:** Definitions, base values, and sources of model parameters.

| Parameter | Default value | Description | Source |
| --- | --- | --- | --- |
| $r_A$ | 12 yr<sup>-1</sup> | Kelp growth rate | |
| $K$ | 4 kg m<sup>-2</sup> A | Kelp carrying capacity | Cavanaugh et al. 2011 |
| $\delta_A$ | 1.1 U<sup>-1</sup>yr<sup>-1</sup> | Urchin grazing rate on kelp | Lauzon-Guay et al. 2009 |
| $\delta_U$ | 11 P<sup>-1</sup>yr<sup>-1</sup> | Predator attack rate on urchins | Tegner and Levin 1983 |
| $a$ | 0.41 UA<sup>-1</sup> | Kelp to urchin conversion | see Appendix S1 |
| $b$ | 0.014 PU<sup>-1</sup> | Urchin to predator conversion | see Appendix S1 |
| $f_c$ | 0.85 | Predator recruitment facilitation | see Appendix S1 |
| $s_A(\tau_K, x)$ | 0.305 (mean) | Winter kelp survival | see Appendix S1 |
| $\sigma_U(\tau_K, x)$ | 1 (mean) | Urchin larval survival stochasticity | see Appendix S1 |
| $\mu_U$ | 0.1 yr<sup>-1</sup> | Urchin natural mortality | Marzloff et al. 2013 |
| $\mu_P$ | 0.05 yr<sup>-1</sup> | Predator natural mortality | Neilson 2011; Hamilton et al. 2011 |
| $F_P$ | 0.19 yr<sup>-1</sup> | Predator harvest rate | Neilson 2011; Hamilton et al. 2011 |
| $\beta$ | 6.7x10<sup>-5</sup> yr<sup>-1</sup>m<sup>-2</sup> | Non-urchin predator resources | |
| $\Delta x$ | 1.5 km | Length of patch along coast | |
| $\psi_U$ | 80 km | Mean urchin dispersal distance | Edmands et al. 1996 |
| $\psi_P$ | 80 km | Mean predator dispersal distance | Waples and Rosenblatt 1987 |
| $\psi_{s_A}$ | 20.8 km | Distance at 0.5 correlation in $s_A$ | see Appendix S1 |
| $\psi_{\sigma_U}$ | 12.0 km | Distance at 0.5 correlation in $\sigma_U$ | see Appendix S1 |

### Model description

Here we describe the model of local community dynamics (Fig. 1a), and in the next sub-sections we describe the 1-patch and spatially explicit formulations of the model (Fig. 1 b, c). Within each patch *x*, our model tracks kelp biomass (algae, *A*_*x*_) and densities of adult consumers (urchins, *U*_*x*_, and predators, *P*_*x*_). We use a semi-discrete (Mailleret and Lemesle 2009) model where kelp growth, herbivory, predation, and harvest mortality occur continuously for one year (*t* ≠ *τ*_*K*_), followed by an annual discrete-time pulse (at *t* = *τ*_*K*_) with storm-induced kelp mortality and settlement of newly produced urchins and predators into local communities.

**Figure 1:**
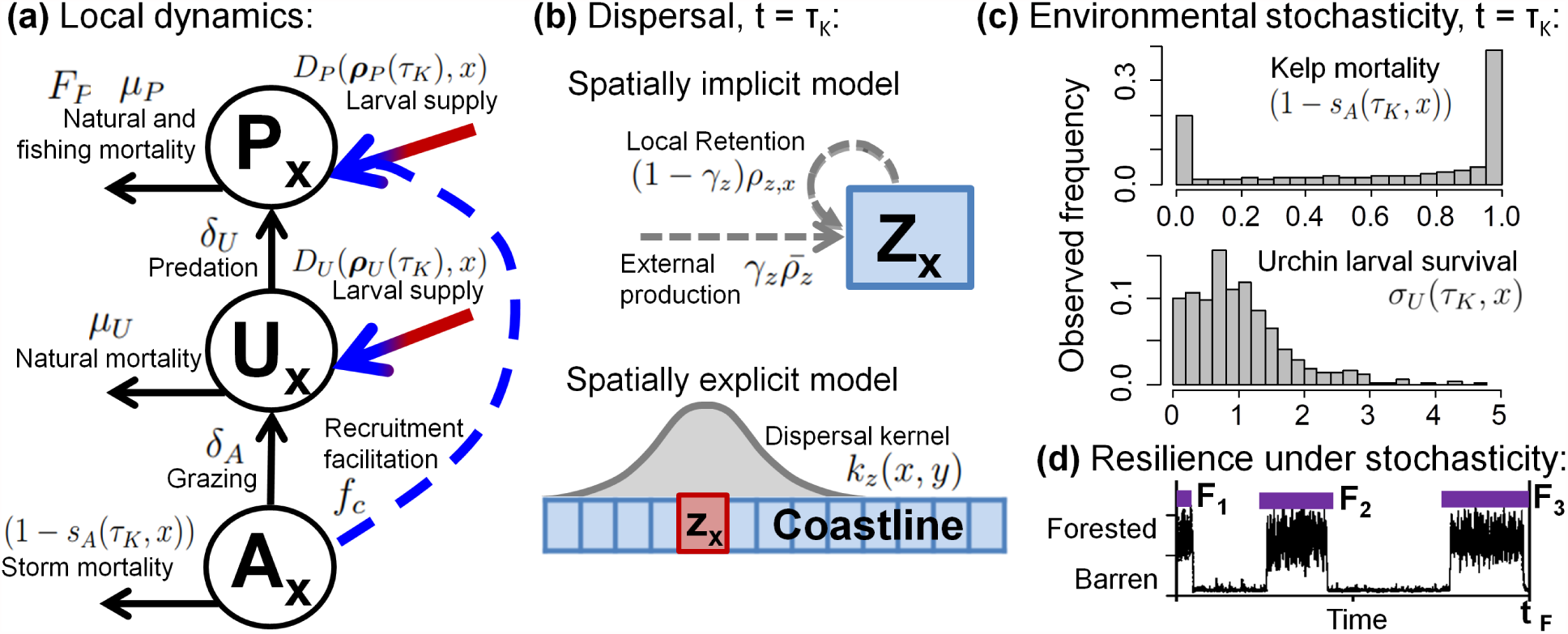
Model schematic of kelp (A), urchin (U), and urchin predator (P) dynamics within a local community x (panel a), denoting losses to consumption and mortality and consumer settlement dynamics at the reproductive pulse *t* = *τ*_*K*_. Arrow colors denote pre-settlement (red), settlement (blue), and post-settlement processes (black). Prior to settlement, larvae of each consumer *z* disperse in both 1-patch and spatially explicit models (b). Kelp mortality and urchin larval survival (relative to average conditions) implemented at the reproductive pulse are determined from empirically observed environmental stochasticity (c). Time series in (d) illustrates two stochastic metrics of resilience used in model analysis: state frequency calculated as 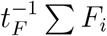 and mean state duration calculated as *n*^−1^ ∑ *F*_*i*_.

Throughout the year, kelp biomass grows logistically at a rate *r*_*A*_ with carrying capacity *K*, and urchins graze kelp at a rate *δ*_*A*_. Urchins and predators experience natural mortality at rates *µ*_*U*_ and *µ*_*P*_, and predators consume urchins at rate *δ*_*U*_ and experience fishing mortality at a rate *F*_*P*_. Thus, the community dynamics within a year (at 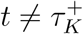) are:

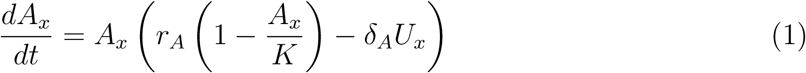

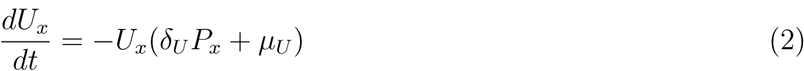

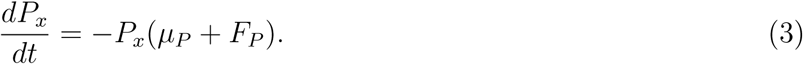

Both urchins and predators experience an annual reproductive pulse at time *τ*_*K*_ (spring-early summer), where larvae produced within each patch disperse between patches. Larval production in each patch is proportional to grazing or predation in the local community integrated over the previous year. Urchins convert consumed kelp into larvae (accounting for larval mortality during development) given the conversion factor *a*. Two energetic sources contribute to predator fecundity: consumption of urchins given the conversion factor *b* and feeding on alternative (non-urchin) prey species *β*, where *β* ≤ *µ*_*P*_ to prevent unbounded predator growth. We multiply this per capita larval production by the number of individuals that live to reproduce in early spring to arrive at the larval production for species *i* in patch *x*, *ρ*_*i,x*_(*τ*_*K*_):

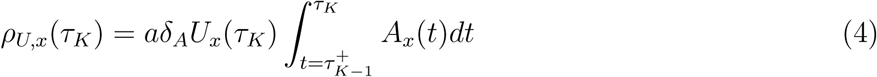

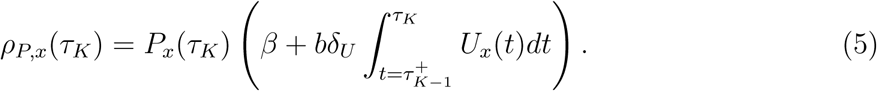

Given larval production in all *n* patches in the vector ***ρ***_*i*_(*τ*_*K*_), we account for dispersal with the function *D*_*i*_(***ρ***_*i*_(*τ*_*K*_), *x*), which depends on the spatial model. Urchin larvae can then experience stochasticity in the number of dispersing larvae that end up in a population *σ*_*U*_ (*τ*_*K*_, *x*), which captures pre-settlement larval survival and movement by water currents (Hart and Scheibling 1988), settlement success dependent on factors such as local cues, and post-settlement predation on juveniles (Schroeter et al. 1996), where our distinction between “pre-settlement”, “settlement”, and “post-settlement” phases follows (Benton and Bowler 2012; Bertness et al. 2014). Because larval survival predominates stochasticity across these processes, from this point we refer to *σ*_*U*_ (*τ*_*K*_, *x*) as stochasticity in larval survival. In preliminary simulations we found that compared to urchin larval survival, stochasticity in predator larval survival had only weak effects on community dynamics because predators experience much lower population turnover, so we ignore it here. To account for recruitment facilitation, successful settlement of predator larvae increases proportionally with local kelp biomass by a factor *f*_*c*_, where 1 − *f*_*c*_ is the baseline predator settlement success in the absence of kelp (Baskett and Salomon 2010). Finally, kelp biomass experiences stochastic survival *s*_*A*_(*τ*_*K*_, *x*) due to storms and limited growth during winter-spring. See the *Model implementation* section below for details on the distributions of *s*_*A*_(*τ*_*K*_, *x*) and *σ*_*U*_ (*τ*_*K*_, *x*) for different model realizations. Thus, the community state just after the reproductive pulse (at 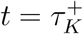) is:

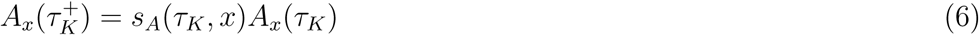

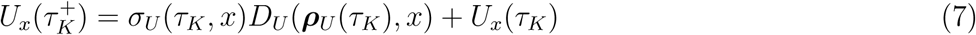

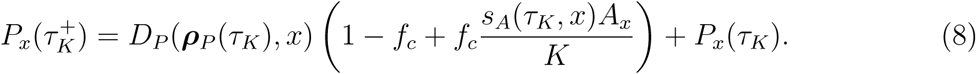

### Spatial dynamics

We start with a 1-patch model to link to existing theory and measurement of resilience as the size of a basin of attraction (see *Model analysis* below). We then extend the framework to a spatially explicit model to test the robustness of our results to greater realism and determine the spatial scale of alternative stable states where relevant. In the 1-patch model (Fig. 1b), we incorporate external larval production as the proportion *γ*_*i*_ of urchin and predator larvae that originate from outside communities. Thus, *γ*_*i*_ = 0 corresponds to demographically closed and *γ*_*i*_ = 1 corresponds to demographically open populations following Johnson 2005. Combining larval production in external areas for species *i* 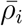 with local retention, dispersal into the local community is

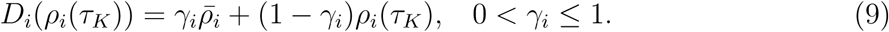

In the spatially explicit model (Fig. 1c), the density of species *i* larvae arriving in patch *x* depends on the dispersal kernel *k*_*i*_(*x, y*), which describes the proportion of larvae in patch *y* dispersing to *x*:

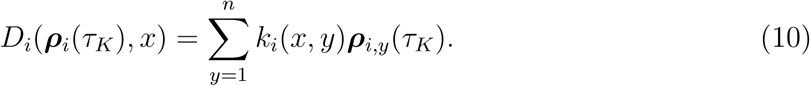

For the discrete dispersal kernel, we integrate a continuous Gaussian distribution with mean 0 and variance 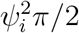 (i.e., mean dispersal distance *ψ*_*i*_ when integrating absolute dispersal distances over all offspring) over the length of coast ∆*d* spanned by patch *x* to calculate

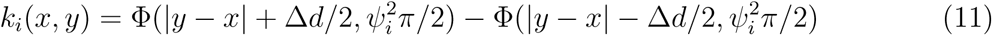

using the Gaussian cumulative density function Φ.

### Model implementation

We numerically simulate model dynamics using R 3.4.3 (R Core Team 2017), solving intrannual dynamics 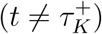, and accounting for dispersal and environmental stochasticity at the end of each year 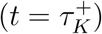. We use a spatial resolution of ∆*d* = 5.5 km (i.e., *n* = 200 patches) to resolve the smallest-scale stochasticity, and analyze simulations over 1000 years following a 3000 year burn-in period to exclude transient dynamics (Appendix S1: Section S3). Given the presence of kelp forests outside our study area, we exclude edge effects by assuming a linear coastline with periodic boundary conditions in *k*_*i*_(*x, y*) (Gouhier et al. 2010), placing organisms that disperse beyond the southern or northern limits of our system into patches on the opposite end.

We parameterize our models based on published studies (Table 1) and monitoring data (Appendix S1: Section S1). In one-patch models, we calculate external larval production 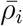 as would occur for 75% kelp forest in all suitable habitat in the system (observed in Castorani et al. 2015; we relax this assumption in Appendix S1: Section S1, Fig. S4). For this we separately determine larval production in each state for closed communities. In the stochastic case we calculate production over a transient period that precedes eventual predator declines to the long-term absorbing state of extinction (Appendix S1: Section S1) that occurs without external larval supply. In deterministic simulations we stochasticity in urchin larval survival (*σ*_*U*_ (*t, x*) = 1) and use average kelp survival (*s*_*A*_ = 0.37). In stochastic simulations we model *s*_*A*_(*τ*_*K*_) as a beta distribution fitted to remote-sensed kelp biomass in the region (31 years, Bell et al. 2015; Fig. 1c) and *σ*_*U*_ (*τ*_*K*_, *x*) as a beta distribution fitted to observed age-1 urchin densities (27 years, 33 sites, Kushner et al. 2013). We then rescale *σ*_*U*_ (*τ*_*K*_, *x*) by its mean so that stochasticity does not affect long-term larval survival (i.e., average *σ*_*U*_ (*τ*_*K*_, *x*) = 1). In the spatial model we account for observed scales of environmental stochasticity (*ψ*_*s*_*A*__, *ψ*_*σ*_*U*__; Appendix S1: Section S1) using multivariate beta distributions, wherein correlations in kelp and urchin larvae survival decline with distance according to a Gaussian distribution (discretized as in eqn. 11).

### Model analysis

In one-patch models, we measure resilience *R* of the kelp forest state, with alternative stable states present for 0 *< R <* 1, and absent for *R* ≈ 1 (indicating 100% forests) and *R* ≈ 0 (indicating 100% barrens). In the deterministic case we quantify R as the proportion of 1000 uniformly spaced initial conditions that result in kelp forests. In stochastic simulations we measure R as the proportion of time local communities spend in a forested state (Scheffer et al. 2015). We also measure the mean duration of forest and barren states, where increasing durations signify increasing resilience of each state (Fig. 1d; Livina, Kwasniok, and Lenton 2010). In all stochastic simulations we first test for multimodality in the distribution of kelp biomass using Hartigans’ Dip test (Hartigan and Hartigan 1985) to see if alternative stable states are present. We then identify each state by fitting mixed skewed-normal distributions to model results, and use these distributions to determine community states in every patch and year (i.e., *A*_*x*_(*τ*_*K*_), *U*_*x*_(*τ*_*K*_)). Note that while predators are largely extinct in patches with lengthy barren states, predator densities in stochastic models generally vary over longer time scales and do not exhibit a clear bimodality.

In spatially explicit simulations, we first test for system-wide alternative stable states by investigating the initial-condition dependence of the system-wide outcome. Specifically, we measure the frequency of forest states, aggregated across space, in simulations starting from either all patches initially forested or nearly barren. In stochastic simulations where disturbances might induce localized state shifts, we quantify the mean spatial extent and duration of forests and barrens over the range of fishing intensities. To verify that our stochastic dynamics reflect the presence of alternative stable states (Scheffer et al. 2015), we run simulations with and without the recruitment facilitation feedback (i.e., *f*_*c*_ = 0.85 and 0) and test for bimodality in kelp biomass. Bimodality with and unimodality without facilitation indicates that feedback-based alternative stable states drive community dynamics, while bimodality in both cases indicates stochasticity as the driver of observed patterns. Additionally, to evaluate whether the scale of disturbance (*vs.* dispersal) determines the scale of alternative stable states, we explore a range of spatial scales in urchin settlement stochasticity 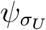.

### Comparison to observed community scales

To test the realism of our spatial model predictions, we compare the spatial scale and durations of alternative states in the model to monitoring data from 83 sites across the Channel Islands (Kushner et al. 2013; PISCO et al. 2011; see Appendix S1: Section S4 for analysis details). Monitoring sites span only kelp habitats where kelp loss arises predominantly from urchin grazing; consequently, we classify each site in every year as forested if adult plants are present (*>* 1 ind. 100m^−2^) or barren otherwise. Due to the limited length of time series (10 to 22 years), most (76%) observations record only partially observed state durations (i.e., start and/or end years unknown). To utilize all data, we compare observations with 10000 equivalently long subsets of model simulations, each starting at a randomly selected patch and year (omitting model transients; see Appendix S1: Section S4 for comparison of only fully observed states). Because data were recorded at individual sites, we calculate the spatial scale of states in both simulations and data as the distance at which Pearson correlation in community state *κ*, calculated over a 5km window moving by 0.5km, declines to zero (with standard error 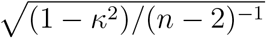 given n comparisons).

## Results

### Presence of alternative stable states under external larval input

We find that the occurrence of alternative stable states decreases with the fraction of external predator larvae but increases with the fraction of external urchin larvae in both the deterministic (Fig. 2a) and stochastic (Fig. 2b-d) implementations of our model. Intuitively, external urchin larval input increases the persistence of barren states by maintaining high settlement when kelp are overgrazed to low levels. External urchin larval input also can reinforce the forest state because, given high predator recruitment facilitation in forests, greater urchin settlement and therefore predator consumption translates into higher predator densities, resulting in more persistent forest states. This outcome entails high urchin population turnover in forest states, as observed in reality (Kenner and Lares 1991). Overall, external urchin larval input maintains the presence of alternative stable states and reduces forest resilience. The magnitude of annual urchin settlement, coupled with fast growth of kelp biomass, also means that pulses of high and low urchin larval survival are the primary source of environmental stochasticity that drives community shifts in our model (Appendix S1: Section S5).

**Figure 2:**
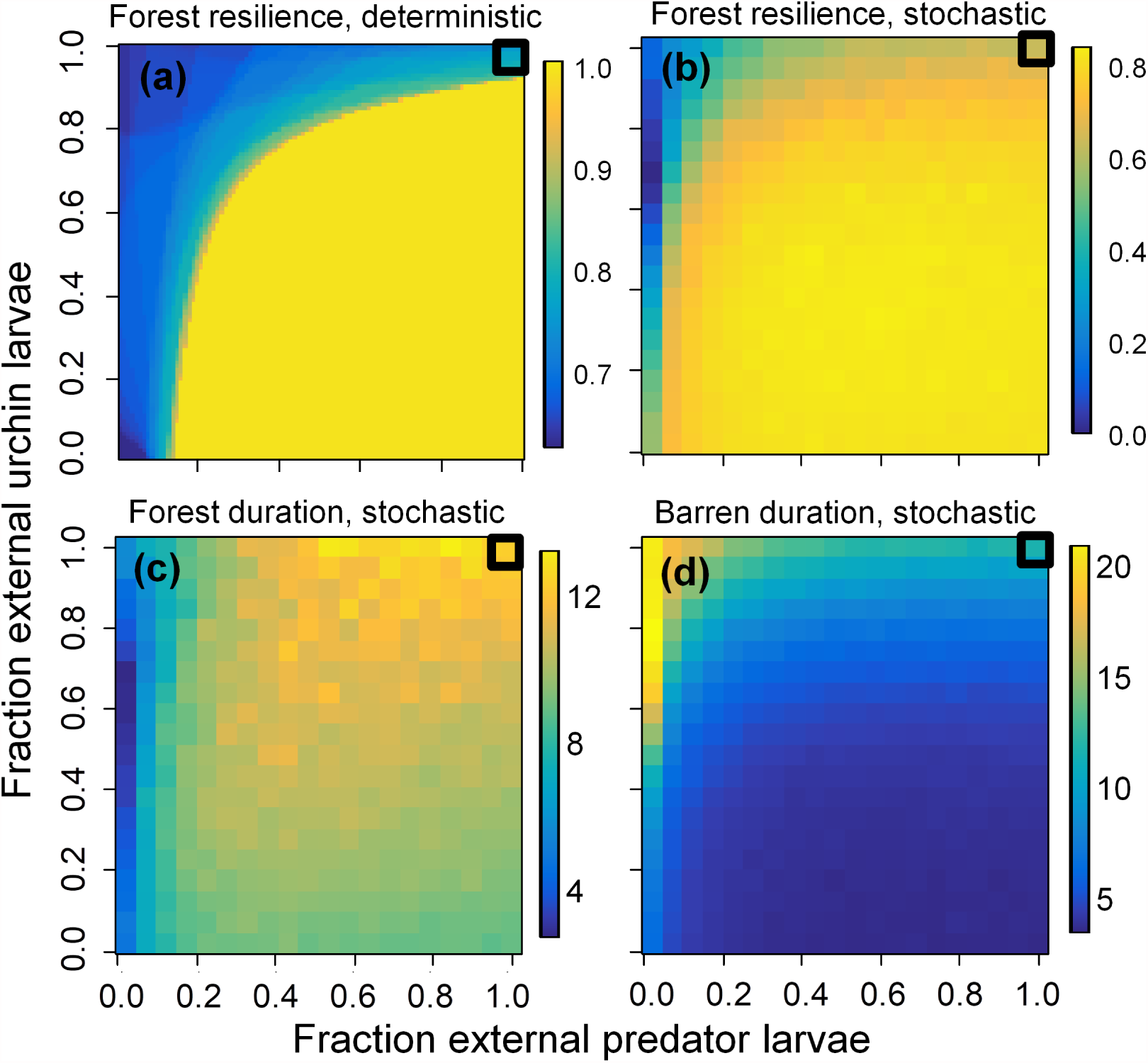
Effects of external predator and urchin larval supply from a partially (75%) forested environment on the presence of alternative stable states (i.e., kelp forest resilience between 0 and 1) without (a) and with (b) environmental stochasticity, and the effects of external larval supply on mean duration of forest (c) and barren (d) states in the stochastic model. For reference, autocorrelation in community dynamics without recruitment facilitation (*f*_*c*_ = 0) declines to zero in 4-5 years. Boxes highlighting larval supply regimes in the top-right of each panel denote the proportion of external larval supply corresponding to our spatially explicit model (dispersal distances *ψ*_*i*_ = 80*km* are approximately equivalent to a fraction of external larvae *γ*_*i*_ = 0.997; see Appendix S1: Fig. S1 for the full relationship between *γ*_*i*_ and *ψ*_*i*_).

In contrast to urchins, the fraction of external predator larvae increases the frequency and duration of kelp forests by increasing total settlement of predator larvae in urchin barrens. The feedback driving alternative stable states (recruitment facilitation) limits this outcome by controlling the number of dispersing individuals that successfully settle within a location. Taken together, external predator larval supply alone leads to loss of alternative stable states (only forests stable), while at high external larval supply in both species, effects of external urchin larval input predominate, maintaining barrens locally even in regions dominated by kelp forests. This presence of alternative stable states under high external larval input requires feedbacks that affect predator settlement, and disappears in a preliminary investigation of post-settlement feedback mechanisms that affect only established individuals (Appendix S1: Section S2, Fig. S5).

### Spatial scales of alternative stable states

Consistent with the 1-patch simulations, our stochastic spatially explicit model reveals forested and barren states at localized but not global scales. While the deterministic spatial model exhibits initial-condition-dependent system-wide alternative stable states at intermediate levels of fishing intensity (the predominant source of predator mortality, Fig. 3a), in the presence of stochasticity the long-term, spatially aggregate ecosystem state is independent of initial conditions (Fig. 3b; Appendix S1: Fig. S7). However, individual patches in the stochastic simulations typically occur in one of two distinct urchin- and kelp-dominated states (87% of patches with *<* 0.02 or *>* 1*kg m*^−2^ kelp, Fig. 4c) that span localized 10-20 km scales and can persist for decades (Fig. 5a, b). This feature reflects alternative stable state dynamics because the stochastic model without the recruitment facilitation feedback exhibits a unimodal state distribution (Fig. 4b, c).

**Figure 3:**
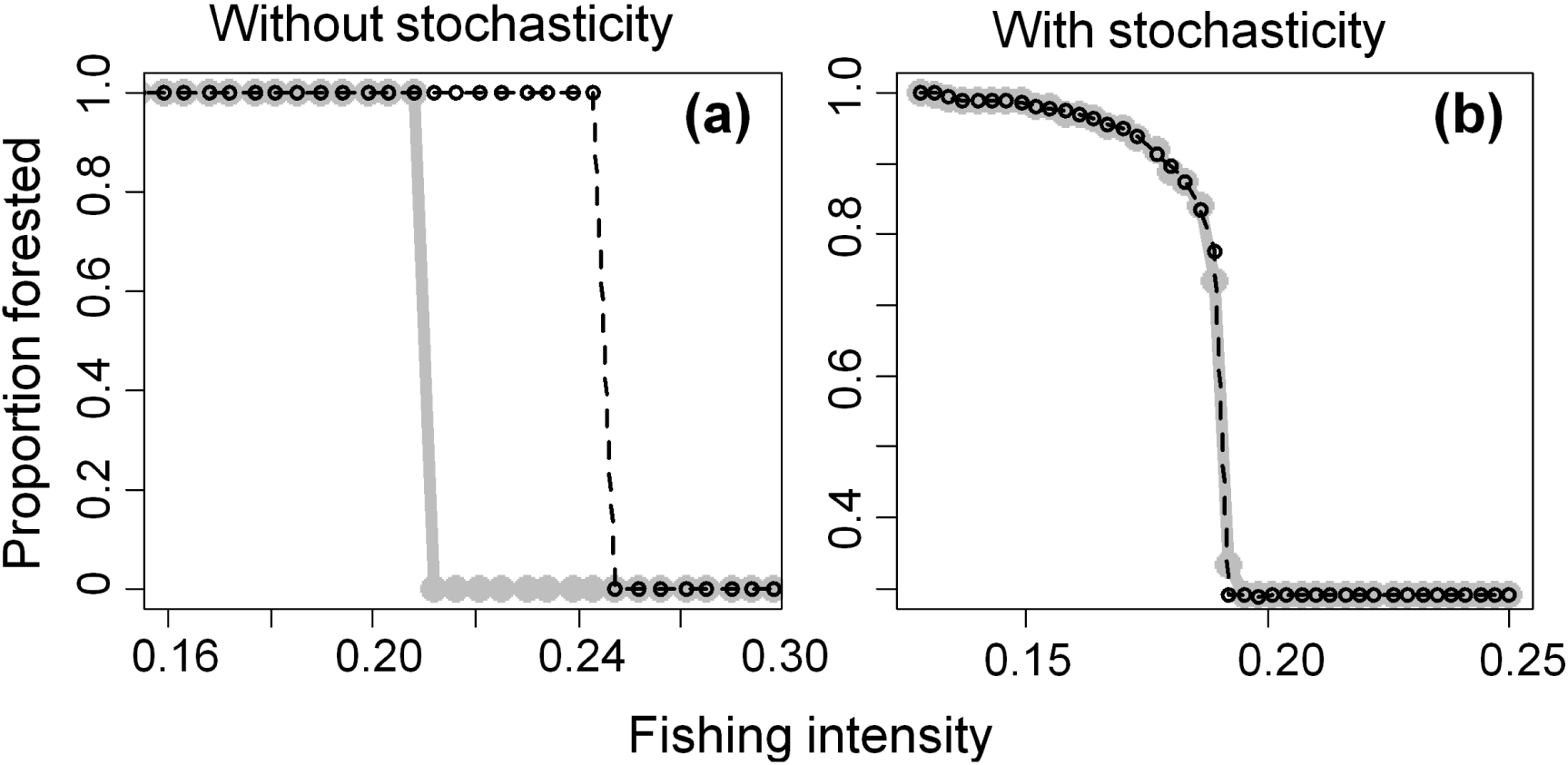
System-wide alternative stable states that occur in the deterministic spatially explicit model (a) disappear in the stochastic model with environmental variation (b). Each plot shows long-term community dynamics of simulations starting from a forested (black, dashed lines) or near-barren (gray, solid lines) state, across levels of fishing intensity, the predominant source of predator mortality.

**Figure 4:**
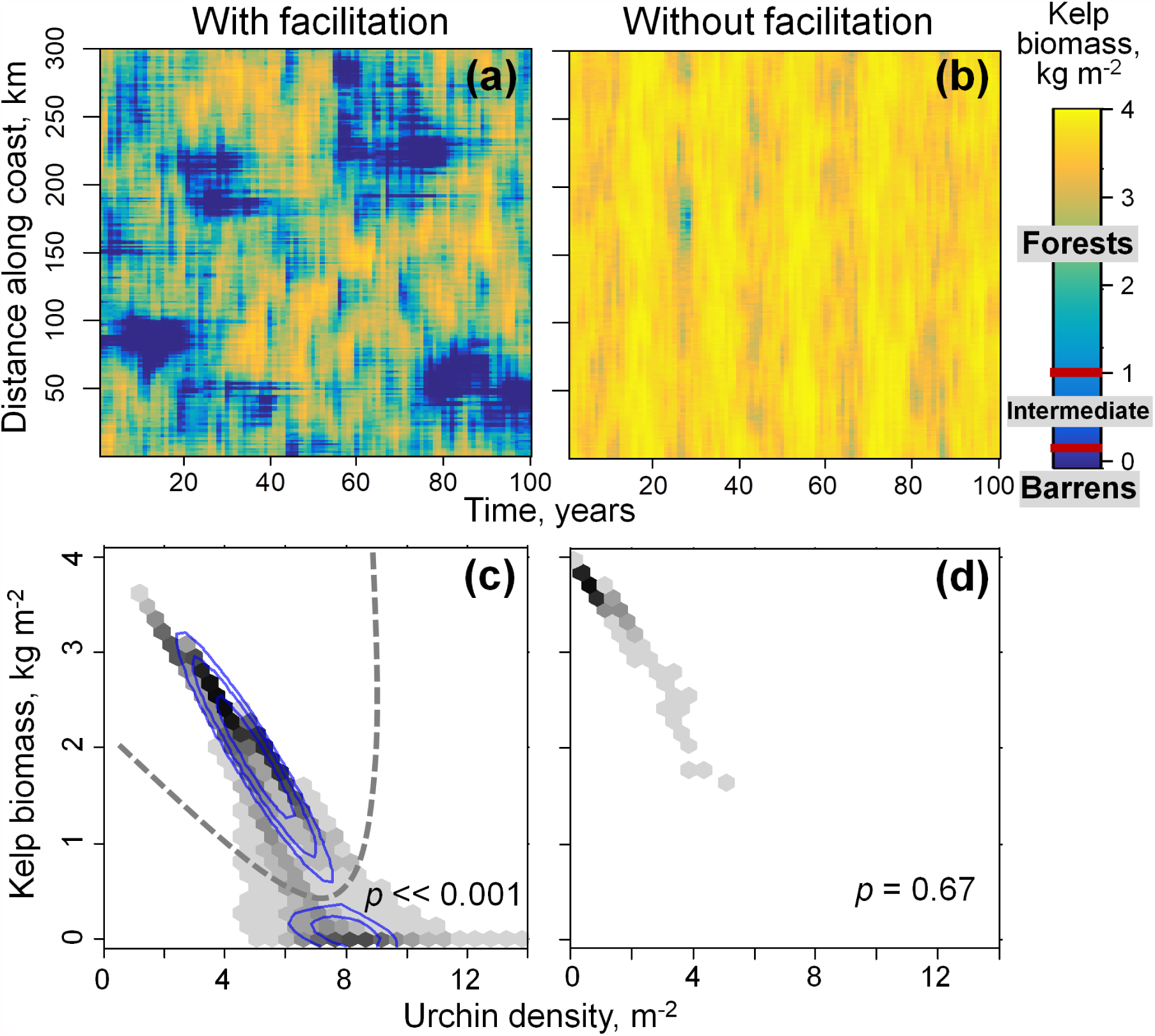
Spatiotemporal kelp forest dynamics (top row) and frequency distributions of community states (bottom row) in our model with (*f*_*c*_ = 0.85, left column) and without (*f*_*c*_ = 0, right column) the recruitment facilitation feedback that produces alternative stable states. (a) and (b) show kelp biomass, with forested states in green-yellow and barren states in dark blue. The time period shown is from after the 3000-year burn in period from a spatially homogeneous kelp forest state. *P*-values from the Dip test in (c,d) indicate whether the distributions of kelp biomass are multimodal (*p <* 0.05), indicating the presence of alternative stable states, or unimodal (*p >* 0.05). In (c) contours denote the best-fit skew-normal bivariate distributions, and dashed line denotes the fitted boundary between forest and barren states.

**Figure 5:**
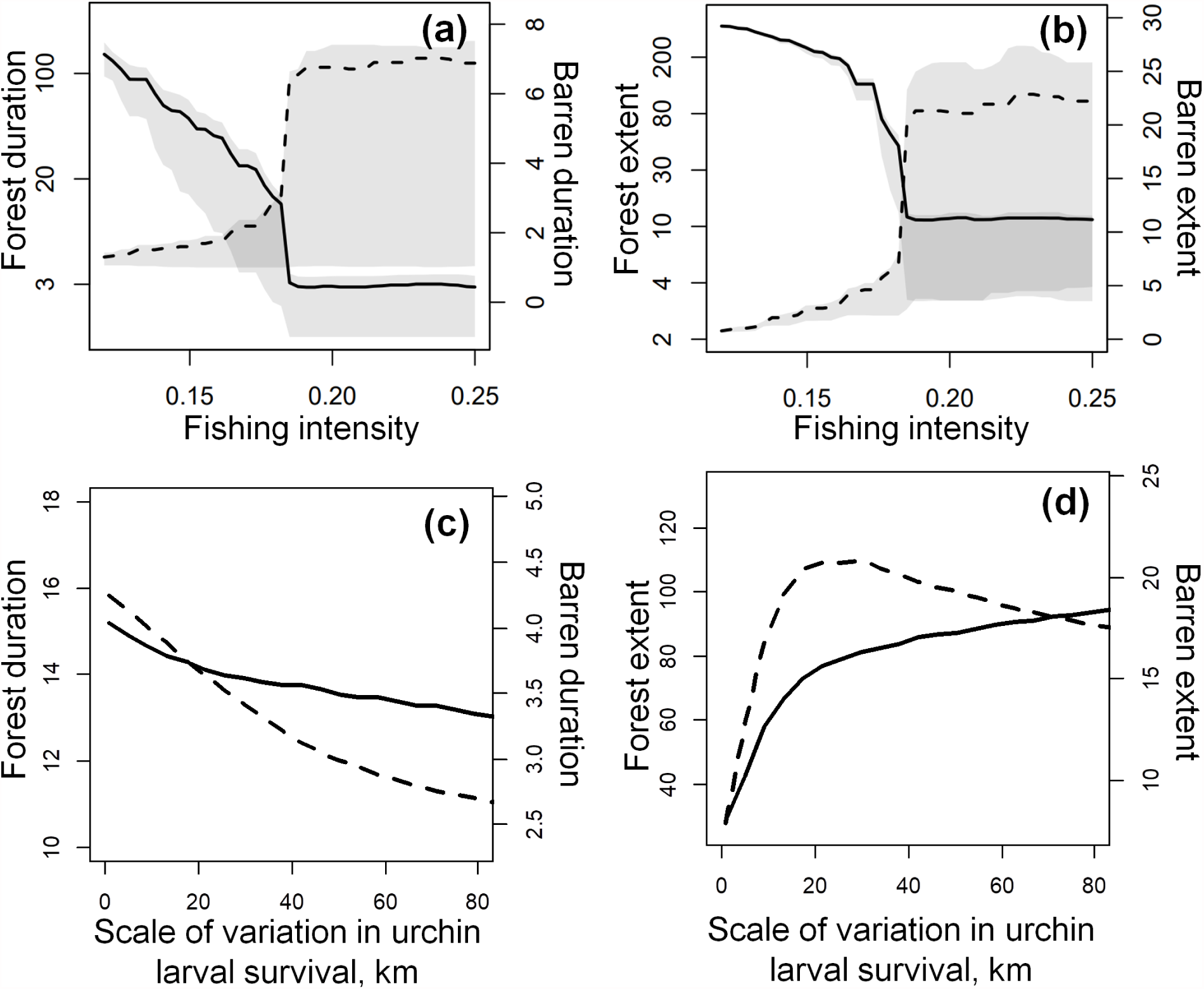
Effects of fishing intensity (which determines forest resilience, a-b) and the spatial scale of variability in urchin larval survival (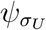, c-d) on the average duration (a, c) and spatial extent (b, d) of kelp forests (solid lines) and urchin barrens (dashed lines). Shaded areas in (a) and (b) denote variation (25th and 75th percentiles) in state duration across patches (a) or spatial extent across years (b). Parameters not varied in each plot (i.e., *F*_*P*_ or 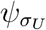) held at default values (Table 1).

Our spatially explicit model also illustrates how the spatial scale of alternative stable states depends on the interaction between the scale of environmental stochasticity and the amount of ecological resilience based on within patch dynamics. By reducing forest resilience, higher fishing increases the frequency and duration of barrens (and vice versa for forests; Fig. 5a, b) as pulses of urchin larvae become more likely to produce barren states. The spatial scale of barrens simultaneously increases because, for a given disturbance level, nearby locations experience greater likelihood of being in a barren state. The spatial scale of urchin barrens peaks at intermediate spatial scales of environmental stochasticity (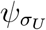; Fig. 5c, d): initial increases in the scale of variation in larval supply increase the scale of both states by synchronizing environmental conditions among nearby communities. Eventual increases in 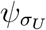 reduce the capacity of urchin dispersal to mitigate stochastic population declines via larvae from areas experiencing favorable environments (i.e., the spatial storage effect). The resulting declines in overall urchin densities and settlement reduce the extent of barrens by reducing their resilience and the persistence of both states.

### Comparison to observed community scales

Duration and spatial scales of alternative states in the spatial model were partially comparable to data (Fig. 6). Overall, distributions of observed forest state durations were not significantly different from model expectations (D=0.07, *P* = 0.61, two-sample Kolmogorov-Smirnov tests here and throughout). However, distributions of barren durations differed significantly (D=0.27, *P* < 0.001), with mean barren durations shorter in simulations compared to data (2.9 *vs.* 4.8 yr; see Discussion for possible explanation). Omitting partially observed states yielded analogous results (Appendix S1: Section S4). The spatial scale of community states was on a similar magnitude in simulations compared to data, albeit substantially larger (15 km *vs.* 30 km, distances where correlation declines to zero in Fig. 6c).

**Figure 6:**
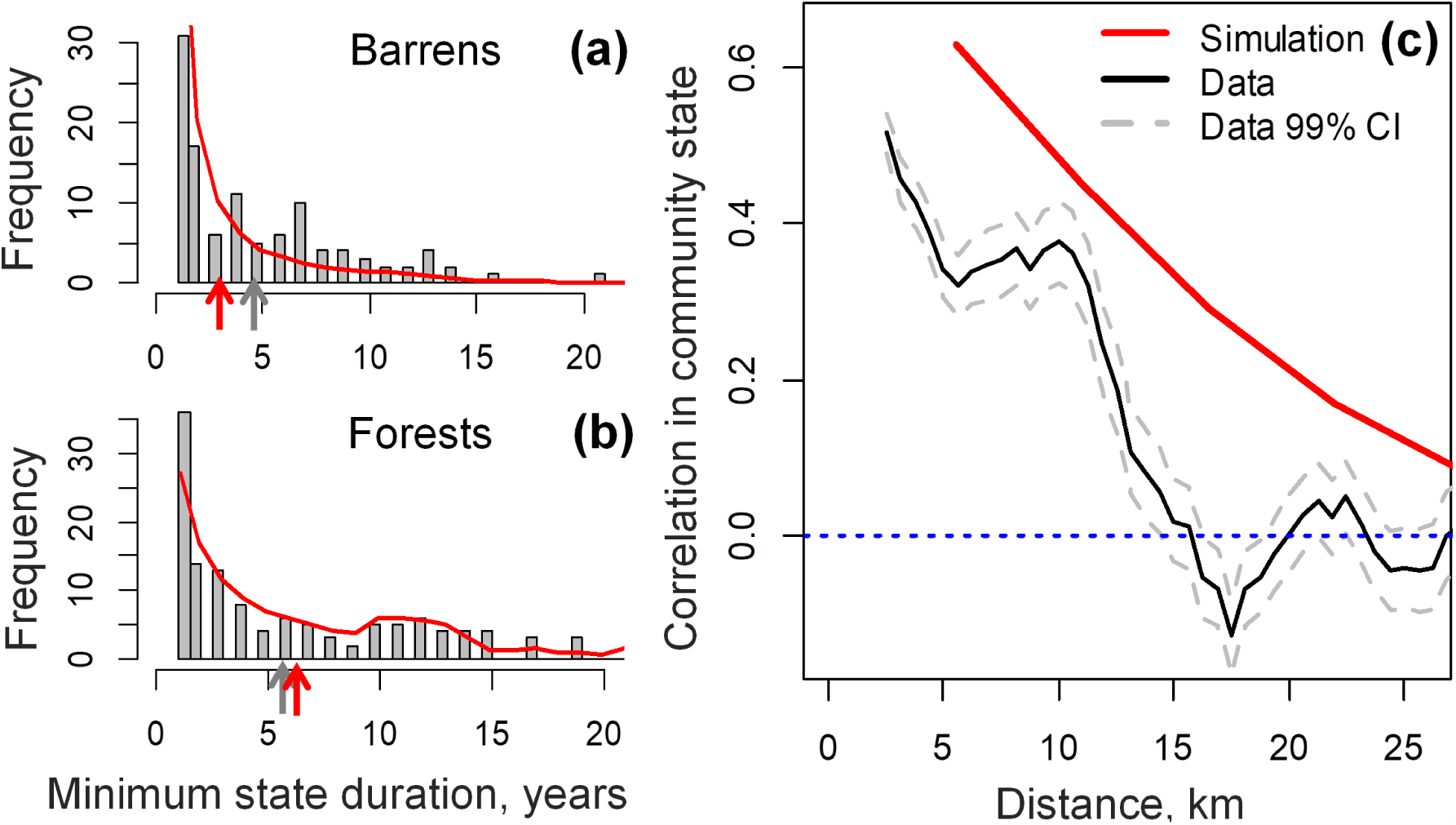
Durations (a, b) and spatial scales (c) of community states in model simulations compared to field data. Minimum durations (i.e., both fully and partly observed states) of barrens (a) and forests (b) given for data (bars, frequencies) and model simulations (red). Spatial autocorrelation of community states in simulations (red) and data (black points) is calculated over a 5km window moving by 0.5km; gray lines denote 99% confidence intervals.

## Discussion

Our results show that alternative stable states can remain relevant in highly interconnected communities (Fig. 2, 4). This relevance of alternative stable states in our model most likely arises because the feedback mechanism maintaining urchin barrens (low predator facilitation) affects the settlement of dispersing predators into local communities but not post-settlement processes (Appendix S1: Section S2, Fig. S5). Specifically, dispersing larvae do not overwhelm settlement feedbacks within patches as might otherwise be intuitively expected because few dispersers successfully establish in the community. Across local communities, the settlement feedback reduces dispersal among communities in distinct states and maintains localized, alternatively stable forest and barren states (Fig. 4). In contrast, preceding spatially explicit studies that predict dispersal-induced synchrony of ecological states over large scales (van Nes and Scheffer 2005; Martin et al. 2015) have focused on post-settlement feedbacks as the drivers of alternative stable states. This suggests that the relevance of alternative stable states in highly interconnected communities might require settlement feedbacks, a hypothesis worth testing in alternate model structures.

Our results show that ecosystem heterogeneity required for alternative stable states to be relevant locally (here, spatially asynchronous stochasticity; Fig. 3) may occur below the scales of dispersal (e.g., 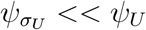 in our model, Fig. 5d). Conversely, in spatially explicit models with post-settlement feedbacks, alternative stable states co-occur among local communities only when stochasticity or habitat patchiness exceed the scales of dispersal (Durrett and Levin 1994; van Nes and Scheffer 2005; Martin et al. 2015). In models with settlement feedbacks similar to ours, fine-scale (0.1-100 m, below the scales of dispersal) patterning of distinct states arises through self-organization when organisms facilitate each other locally and inhibit growth at larger scales (Rietkerk and Van de Koppel 2008). Our findings show the potential for analogous self-organization mechanisms to structure ecosystems on large (e.g., 10-20 km) spatial scales. Taken together, multiple sources of ecosystem heterogeneity can induce alternative stable states at local scales.

We also find that the spatial and temporal scales of each stable state increase with its local-scale basin of attraction and the scale of environmental variation, which outweigh the role of dispersal scale. The attraction basin size of a given state (e.g., forests) depends on localized dynamics (here, fishing intensity, Fig. 5 a, b) and represents its ecological resilience. Our results integrate previous stochastic and spatially explicit models that have separately emphasized how greater resilience increases the persistence (Livina et al. 2010) and spatial scale (Bel et al. 2012) of alternative stable states. The spatial scales of alternative stable states can also increase with the spatial extent of environmental variation via environmentally induced synchrony (Fig. 5 c, d; the Moran effect, Gouhier et al. 2010) when disturbances are well below the scale of dispersal. However, larger-scale disturbances can decrease the spatial extent of both states by reducing the role of dispersal and weakening dispersal-induced synchrony. Analogous dynamics occur in stochastic population models with single equilibria (Kendall et al. 2000), highlighting how extensive research on population synchrony might inform the spatial scales of alternative stable ecosystem states.

### Systems where alternative stable states might be relevant locally

The settlement feedbacks expected to be necessary for the local relevance of alternative stable states in our model are likely to occur in many taxa. In general, plants and numerous marine species disperse during seed or larval stages, which often are inferior competitors compared to individuals already established in the community or rely on facilitating species for survival (reviewed in Caley et al. 1996; Bruno et al. 2003; Clobert et al. 2009). Additionally, actively moving adult animals and aquatic larvae avoid settling in areas of intense competition, high predation risk, or unfavorable conditions (Morgan et al. 1996; Benton and Bowler 2012). Thus, settlement feedbacks can arise from behavior during or high mortality immediately after settlement that prevents dispersers from establishing in the community.

Observed settlement feedbacks might in turn underlie localized alternative stable states in a variety of ecosystems beyond temperate rocky reefs. For tropical reefs where coral larvae cannot settle or survive in algae-dominated areas, alternative stable coral and macroalgal states occur in spatially explicit models on the scale of individual reefs (ca. 500 m^2^) despite high levels of external larval supply (Mumby et al. 2007). Additionally, herbivorous fish that reinforce coral-dominated states preferentially settle in areas of high coral cover (which offer predation refugia) when migrating from nursery habitats (Dixson et al. 2014). Consequently, distinct states might co-occur among local reefs, especially when large-scale disturbances have reef-specific impacts due to long-term, localized environmental differences (e.g., areas with protected *versus* fished herbivores; Mumby et al. 2013). In terrestrial systems where competitively inferior grasses impede tree seedling survival by amplifying fire severity, empirical studies find mounting evidence for alternative stable savanna and forest states (Staver et al. 2011). Consistent with our results, spatially explicit models predict that these states co-occur locally below (and independently of) the scale of tree dispersal (Schertzer et al. 2015) when fires propagate heterogeneously across landscapes. Numerous studies also document localized alternative stable states that arise as fine-scale patterns in systems such as mussel beds, wetlands, and arid ecosystems when established individuals deplete resources or displace growth-inhibiting factors (e.g., snow fields in alpine forests; reviewed in Rietkerk and Van de Koppel 2008). Given that both processes locally inhibit settlement of new individuals, our results imply that the scale of such patterns in general depends on the scales of resource depletion or ecosystem engineering rather than the scale of dispersal.

### Empirical evidence and tests for localized alternative stable states

The localized spatial and temporal scales of alternative stable states predicted here are in line with empirical observations of temperate and tropical rocky reef systems. The duration of kelp forests in our model match observed community scales in the Channel Islands. The greater duration of barrens observed compared to expectations might be due to lower urchin predator densities in the northern Channel Islands compared to the modeled region, which arise partly due to lower body growth or higher mortality (Caselle et al. 2011). The smaller spatial scale of community states in the data compared to expectations likely arises from spatial heterogeneity in rocky reef habitat availability not accounted for here (see *Model robustness* below for details). Worldwide, studies on temperate rocky reefs often find evidence for kelp- and urchin-dominated states (analogous to Fig. 3a) even when sampling at very fine scales (5 m^2^, Ling et al. 2015). Among 51 transitions between these states documented worldwide, two-thirds occurred at < 100 km spatial scales (Filbee-Dexter and Scheibling 2014). These scales are likely at or below the mean dispersal distance of urchins and their predators given the species’ long pelagic larval duration and strong, large-scale genetic exchange (100-300 km; Waples and Rosenblatt 1987; Edmands et al. 1996). Similarly, in tropical coral reefs Ninio et al. (2000) found long-term shifts to algal-dominated regimes that were spatially localized (ca. 10 km) and asynchronous at larger (ca. 100 km) scales.

The spatial scales of alternative stable states found here, while local, still exceed the extent of feasible manipulative experiments and many ecological surveys, which impedes efforts to detect this phenomenon. The two most commonly used approaches to detect distinct states under the same environmental conditions involve demonstrating either (a) a lack of recovery of an original, stable state following experimental manipulations or natural disturbances or (b) a bimodality in the distribution of ecological states observed across locations in field surveys (e.g., Fig. 4c; reviewed in Mumby et al. 2013; Scheffer et al. 2015). We highlight that both features occur only when the spatial extent of experiments or surveys is greater than the spatial scale of alternative stable states; at finer scales, detection efforts might fail because dispersal or long-distance interactions rapidly reverse the effects of manipulations or disturbances, maintaining a single community state (Petraitis 2013). One approach towards resolving the presence and scale of local alternative stable states can be to pair spatially explicit dynamical models with field data (Scheffer et al. 2001). In particular, model selection given observed data can quantify the extent to which observed spatiotemporal patterns are consistent with expectations for multiple types of hypothesized feedback loops, as compared to solely the effects of environmental heterogeneity known to occur in a given system.

### Model robustness

For tractability, we omitted several factors that might affect the feedback loops that drive the relevance of alternative stable states under external larval input. One factor that might erode the recruitment facilitation feedback maintaining urchin barrens in our model is strong predation on urchins by species less reliant on the presence of kelp. Alternatively, several feedbacks omitted here might reinforce urchin barrens. First, changes in urchin foraging behavior not modeled here can lead to intensified grazing at low predator densities (Cowen 1983) or at low levels of kelp biomass and therefore reduced supply of drift kelp (Ebeling et al. 1985; Harrold and Reed 1985). Second, urchins might also experience recruitment facilitation in barrens due to crustose coralline algae, which might compete with kelp (Baskett and Salomon 2010). Additionally, predators preferentially avoid consuming starving urchins in barren areas (Eurich et al. 2014). All of these mechanisms could increase urchin densities or grazing at low kelp biomass. Acting together, multiple feedbacks can produce alternative stable states even when individually each mechanism is insufficient to do so (van de Leemput et al. 2016). The strength of most feedbacks posited to drive alternative kelp and barren states, including our feedback of recruitment facilitation, is often estimated in only a few small-scale, system-specific studies (Ling et al. 2015). Nevertheless, our qualitative results are not sensitive to the level of facilitation (Appendix S1: Section S6), and this mechanism is consistent with greatly reduced predator densities in barrens versus kelp forests in our system (Graham 2004).

Our model also makes several simplifying assumptions that might quantitatively affect the extent and duration of alternative stable states. For example, we ignore the fact that juvenile and adult predators (particularly lobsters) can potentially move over long distances and, unlike larvae, might settle and survive even in barren areas with low kelp cover. Our model also does not account for kelp mortality during extreme warm water events, which can span much larger spatial scales than storm-driven kelp loss (Dayton et al. 1999). Additionally, we parameterize stochasticity in urchin larval survival based on data from the Channel Islands, which experience stronger oceanographic variability compared to other California rocky reefs. Together, increased role of dispersal by adult predator movement, large-scale kelp mortality events, or lower intensity of localized stochasticity might produce larger-scale community states than predicted here. Alternatively, our model might over-estimate these scales due to limited dispersal among communities separated by areas lacking kelp habitats, particularly between islands (5-50 km, Cavanaugh et al. 2013). Other forms of local environmental heterogeneity not modeled here, such as heterogeneity in fishing intensity due to distance from ports or the presence of marine protected areas (Hamilton et al. 2011) and variation in kelp growth conditions might also make forests and barrens manifest within more localized areas of our system.

### Management implications

We find that increasing stress applied to the overall ecosystem (here, fishing) can induce localized socioeconomically undesired states (here, urchin barrens) that gradually increase in frequency, spatial extent, and duration (Fig. 5a, b). Our results suggest that while spatially synchronous state shifts expected in interconnected systems (Hughes et al. 2005, 2013) may be unlikely when settlement feedbacks reduce the role of dispersal (Fig. 3b), collapsed states could span 10-30 km of the coastline and persist for decades under intense exploitation (here, fishing). Where disturbances induce localized undesired states, local management efforts can increase the probability of recovery by increasing the attraction basin size of desired states, for example by reducing predator harvest (i.e., by implementing ecologically sustainable yield *sensu* Zabel et al. 2003). Alternative management approaches to restoring desired states include manipulating species densities, for example by reducing urchin densities in kelp forests (House et al. 2017). However, urchin harvest is practical only at small spatial scales such that maintenance of high potential for urchin larval supply from outside areas can limit efficacy. Furthermore, reversal of barren states may require persistent culling over long periods required for predator recovery.

The persistence of undesired barren states despite high predator larval supply from pristine, forested areas found here (Fig. 2) also suggests that settlement feedbacks can limit the role of rescue effects provided by protected areas in spatially explicit management contexts. However, the overall outcome of reserve-based management will depend both on the magnitude of rescue effects and how reserves alter the attraction basin of desired states in exploited areas. For example, Barnett and Baskett (2015) found that marine protected areas increase attraction basins of states dominated by fished predators in both harvested and protected areas compared to equivalent spatially uniform harvest in a model with well-mixed larval pools. Given the role of heterogeneity in maintaining localized alternative stable states in our spatially explicit model, the next step for understanding reserve effects on resilience in exploited areas are spatially explicit studies that account for the spatial scales of reserves, disturbance, and dispersal.

### Conclusions

We expand existing resilience theory by showing that alternative stable states can occur locally below the scales of dispersal, likely due to feedbacks affecting settlement. This can explain observed community scales on temperate rocky reefs and how alternative states commonly co-occur in highly interconnected ecosystems (Scheffer and van Nes 2004; van de Leemput et al. 2015). In the wide array of marine and terrestrial systems where settlement feedbacks occur, this suggests that anthropogenic impacts such as harvest and grazing can produce increasingly large and persistent undesired states rather than sudden ecosystem-wide collapses.

## Acknowledgments

We would like to thank Robert Dunn, Tom Bell, David Kushner, Jennifer Caselle, Katie Davis, Nicholas Shears, Daniel Reed, Max Castorani, and the Santa Barbara Long Term Ecological Research for biological insights and providing data. We also thank Alan Hastings, Sebastian Schreiber, Jay Stachowicz, and two anonymous reviewers for feedback that greatly improved the manuscript. Funding for this project was provided by a National Science Foundation Graduate Research Fellowship to VAK.

## Appendix S1

### S1 Model parameterization

#### S1.1 Deterministic parameters

Here, we describe the derivation of parameters in Table 1 not taken directly from the literature. Predation and grazing rates, which depend on prey encounter rates and therefore prey per unit area, are inversely proportional to area. Thus, we first adjust consumption rates for differences in spatial scale of our model (i.e., 1 m^2^) and studies estimating these rates. Second, we convert these values to per capita rates by adjusting the published, biomass-based predation and grazing rates for the biomass of each species. We determine per capita biomass based on published biomass-size relations and mean size (urchins: 0.04 kg, Kenner et al. 1992; sheephead: 0.445 kg, Hamilton and Caselle 2015; lobsters: 0.57 kg, Nielson 2011). Finally, we adjust predator attack rate to account for the fact that (1) urchins constitute a fraction of predator diets (30-40%, Hamilton and Caselle 2015; Winget 1968) and (2) only a fraction of predators are sufficiently large to consume urchins (75%, Hamilton and Caselle 2015). Note that sheephead and spiny lobsters experience similar total mortality rates (sheephead: 0.22 yr^−1^, Hamilton et al. 2011; lobsters: 0.17 yr^−1^, Neilson 2011), and can attain similar mean individual mass (see above), densities in kelp forests (sheephead: 0.02 m^−2^, lobsters: 0.025 m^−2^; Caselle et al. 2018), and estimated urchin consumption rates (sheephead *δ*_*U*_ = 0.013, lobsters *δ*_*U*_ = 0.017); therefore, we use parameter values averaged over both species to represent the predator guild.

Given the long duration of urchin and predator larval stages (30-60 days; Edmands et al. 1996; Waples and Rosenblatt 1987), oceanographic models of our system estimate high mean dispersal distances (100 km; Siegel et al. 2003). However, pelagic larval duration can overestimate dispersal distances due to the presence of coastal boundary layers; consequently, we adjust these estimates using a relation derived from oceanographic models of our system (80%, Nickols et al. 2015). To illustrate the connection between the 1-patch and spatially explicit models (Fig. 2, black boxes), we calculate the fraction of external larvae *γ*_*i*_ in a single patch given a consumer dispersal distance *ψ*_*i*_ based on eqn. 11 with *γ*_*i*_ = 1 − *k*_*i*_(0, 0) (Fig. S1). Given highly uncertain larval survival during dispersal, directly estimating the conversion of consumed biomass into production of urchin and predator larvae surviving to settlement is not possible. Instead, we fit these parameter values such that the relative abundance of newly settled urchins and predators in our deterministic model at the kelp forest equilibrium (i.e, 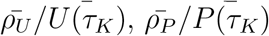) matches observed frequencies of age-1 urchins and predators in our system (urchins: Kenner 1991, 1992; Behrens and Lafferty 2004; sheephead: Caselle et al. 2011; lobsters: Nielson 2011). This ensures that our model captures realistic levels of population turnover in each species. Finally, we estimate the magnitude of recruitment facilitation from the fact that both sheephead and spiny lobsters are 80-90% less common in long-term (i.e., decades) urchin barrens compared to kelp forests (Graham 2004).

**Figure S1:**
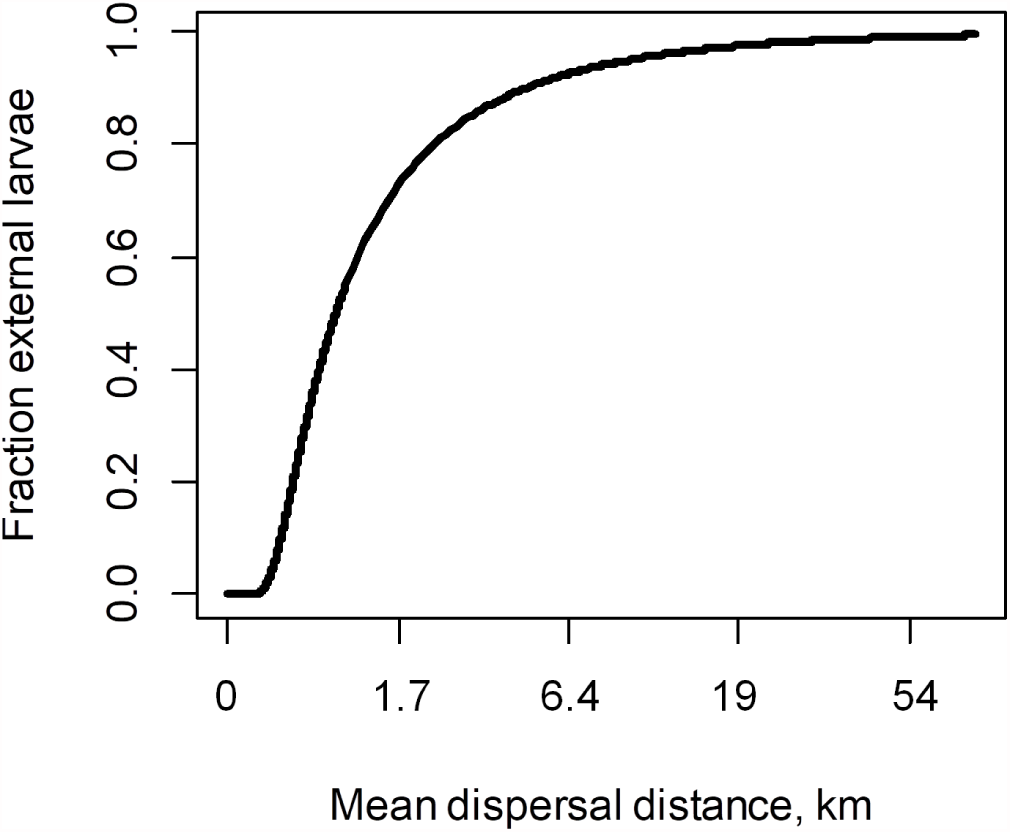
Relationship between mean consumer dispersal distance *ψ*_*i*_ and the fraction of external consumer larvae *γ*_*i*_ in a local community spanning 1.5km of the coastline in our spatially explicit model.

#### S1.2 Environmental stochasticity

For each location and year, we calculate kelp survival as the fraction of maximum kelp biomass observed over the preceding year that remains in the winter and spring (Cavanaugh et al. 2011). We measure *σ*_*U*_ (*τ*_*K*_, *x*) as variation in the density of age-1 (22 − 32 mm test diameter, Kenner 1991) urchins, scaled to the site-level average densities to remove site-specific differences in overall density. To fit a beta distribution to observed distributions of kelp survival (*s*_*A*_(*τ*_*K*_, *x*), Fig. S2a) and density of age-1 urchins (*σ*_*U*_ (*τ*_*K*_, *x*), Fig. S2b) we use the Improved Stochastic Ranking Evolution Strategy (Johnson; Runarsson and Yao 2005). For each parameter evaluation, we compute an empirical cumulative distribution function (ECDF) of both the resulting beta distribution and observed *s*_*A*_ or *σ*_*U*_ levels. Using each ECDF we determine an empirical cumulative distribution (ECD) over 1000 values ranging across observed *s*_*A*_ or *σ*_*U*_ levels and evaluate model fit as the sum of squared differences between the fitted and observed ECDs. Note that we rescale the beta distribution of *σ*_*U*_ (*τ*_*K*_, *x*) fitted and used in our analysis to have a mean of one. In all cases, we complete model fits when relative change in fit of successive parameter sets falls below 0.005 (Fig. S2c, d). This produces model fits *s*_*A*_(*τ*_*K*_, *x*) *∼ Beta*(0.17, 0.3) and *σ*_*U*_ (*τ*_*K*_, *x*) *∼ Beta*(1.8, 20).

To determine the spatial scales of variation in kelp survival 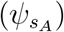 and urchin larval survival 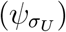, we first calculate the temporal correlation of each factor across pairs of monitoring sites. For 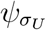 we use sites on the San Miguel, Santa Rosa, and Santa Cruz islands with at least 15 years of data (Kushner et al. 2013). Additionally, we exclude comparisons of *σ*_*U*_ between sites situated on opposite (north versus south) shores of the Channel Islands, which experience an exceptional gradient in oceanographic conditions not present elsewhere in our system. For each factor, we then summarized spatial synchrony by relating correlation and log(1+distance) using least squares, and determined 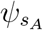 and 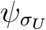 as the distance at which correlation declined below 0.5 (Fig. S2e, f).

**Figure S2:**
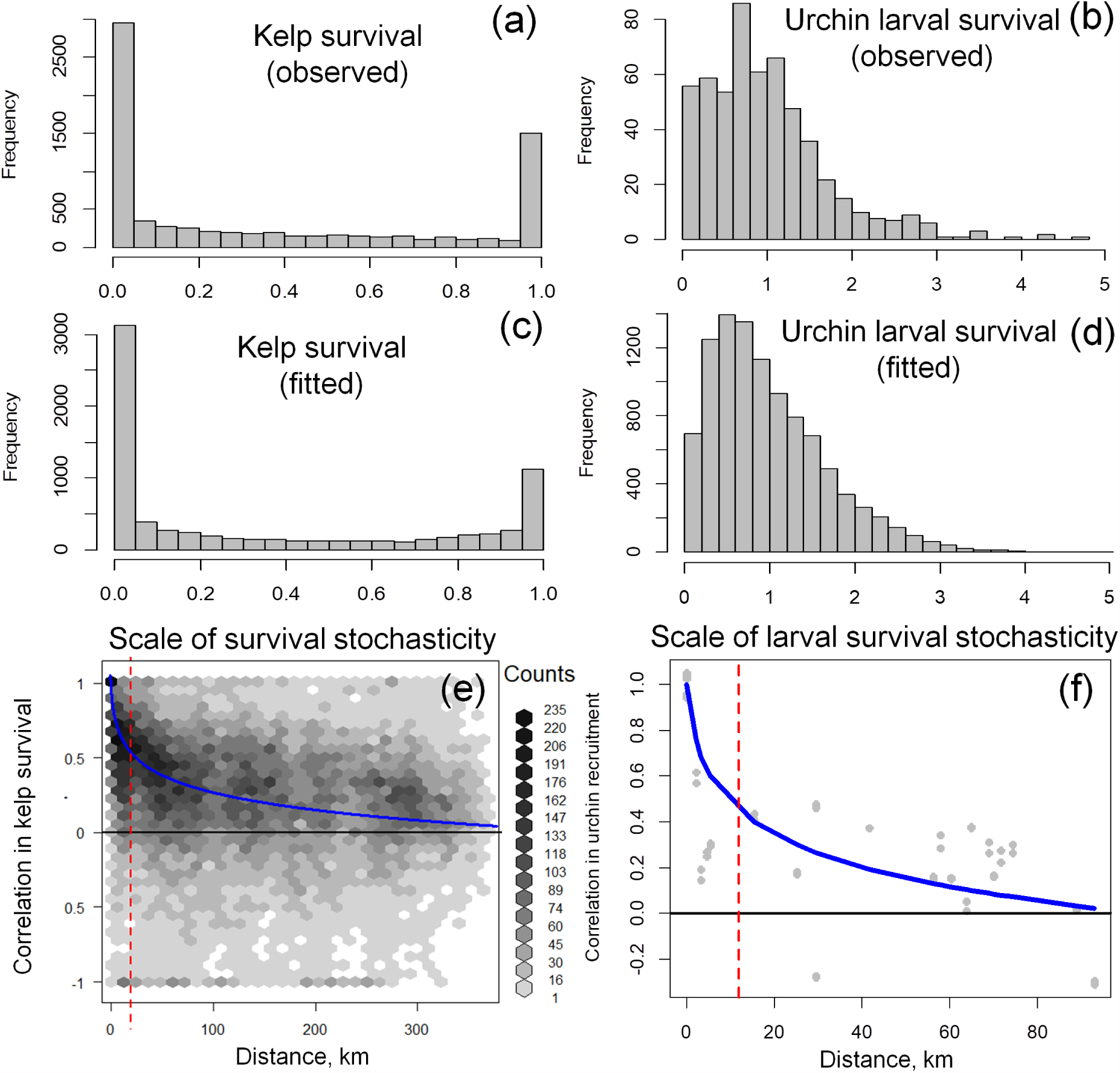
Observed distributions (a, b), fitted distributions (c, d), and observed spatial correlation (e, f) in kelp survival (left panels) and urchin larval survival (right panels). Blue lines in (e, f) show fitted correlation-log distance relationships, and vertical red lines denote the spatial scale of each disturbance 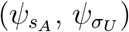 used in spatially explicit simulations.

To determine external larval supply 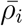 in stochastic one-patch models, we simulate communities initially in a kelp forest state and measure larval production in forested and barren states over a transient period. This period (years 25 to 100) excludes a strong effect of initial conditions and precedes eventual predator extinction, the eventual absorbing state given stochasticity and a lack of external larval supply. Communities within this period exhibit both forested and barren states, and overall maintain persistent predator populations (Fig. S3). Net external larval supply is then a combination of larval production in forested areas (75% in our base model) and barren areas characterized by low urchin fecundity (25% in our base model).

**Figure S3:**
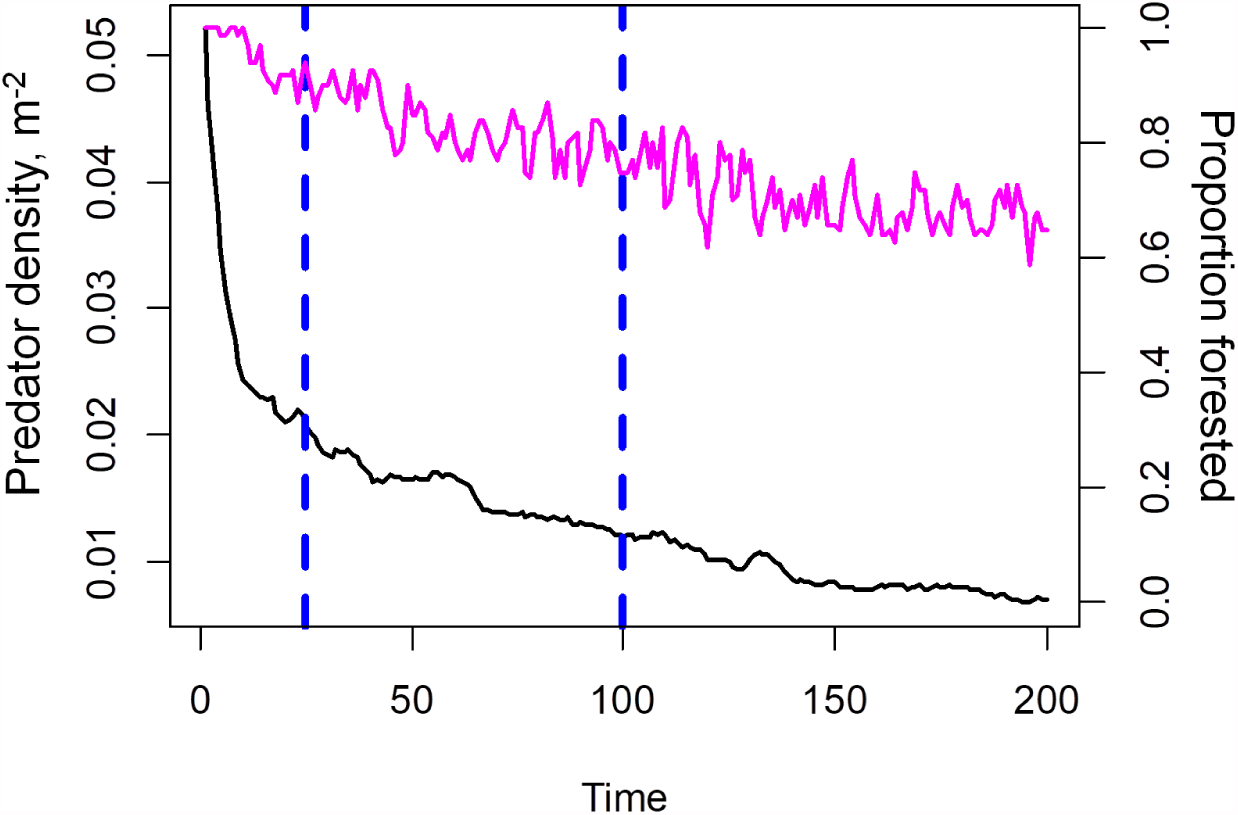
Mean predator abundance (black) and frequency of forested states (pink) across 100 replicate simulations during a transient period (denoted by vertical dashed lines) used to determine urchin and predator larval production in the one-patch models.

As the proportion of outside areas in barren states (modeled implicitly by 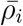) increases, external larval supply affects kelp forest resilience by producing a net export of larvae. Compared to fully forested external conditions (Fig. S4), high external urchin larval supply to partially (25%) barren areas in our main analysis (Fig. 2b-d) results in increased resilience and duration of forest states while decreasing the persistence of urchin barrens. This happens through a net export of urchin larvae when local communities are in a forested state and outside areas include unproductive barren states with low urchin fecundity.

**Figure S4:**
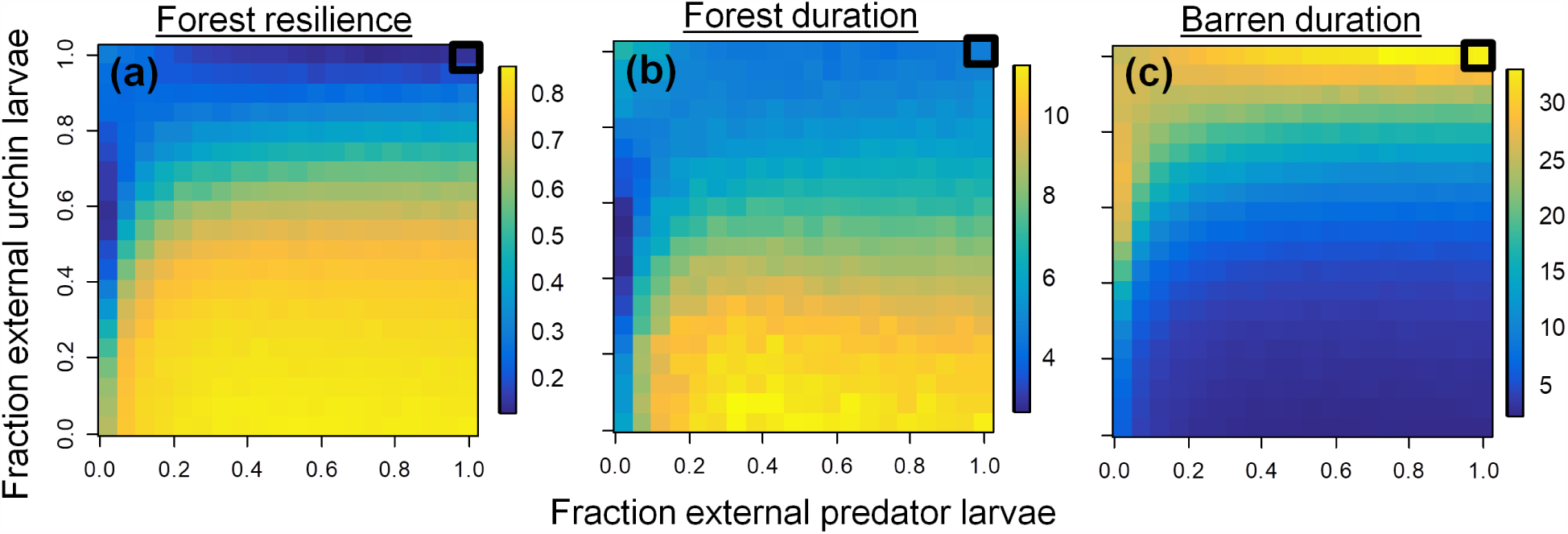
Effects of external predator and urchin larval supply on the presence of alternative stable states (frequency in a and mean state duration in b and c) when external input represents outside areas completely in a kelp forest state. All parameters are as in Table 1.

## S2 Effects of external larval supply with settlement vs. post-settlement feedbacks

Here, we compare how the presence of alternative stable states within a local community with external larval supply depends on the presence of settlement versus post-settlement feedbacks. Whereas our modeled feedback of recruitment facilitation occurs at settlement, models predicting a high likelihood of alternative stable states at system-wide scales in interconnected ecosystems assume post-settlement feedbacks (van Nes and Scheffer 2005). To model this scenario, we modify our one-patch model (eqn. 8 and 9) such that kelp biomass affects local fecundity of established predators but not settlement of arriving larvae. Predator densities at the reproductive pulse then include facilitation only in local production:

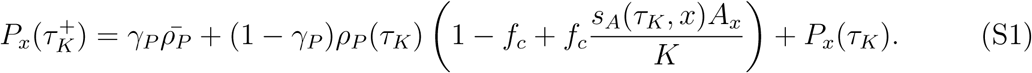

Note that this approach is an initial investigation into how different model structures affect the outcome. Definite establishment of our hypothesis that feedbacks at (rather than after) settlement drive the potential for localized alternative stable states would require a more thorough investigation of multiple model structures, such as feedbacks affecting adult growth and survival, that are beyond th scope of this paper.

In the deterministic analysis of closed communities, models with settlement and post-settlement feedback loops produce identical bifurcation structures in response to fishing intensity (Fig. S5a, d). Consistent with our single-patch stochastic and deterministic (Fig. 2) analyses of external larval supply, alternative stable states occur under both low and high external larval supply under settlement feedbacks (Fig. S5b, c). In contrast, under post-settlement feedback loops alternative stable states disappear beyond very low levels of external larval supply as predator input destabilizes the urchin barren state (Fig. S5e).

**Figure S5:**
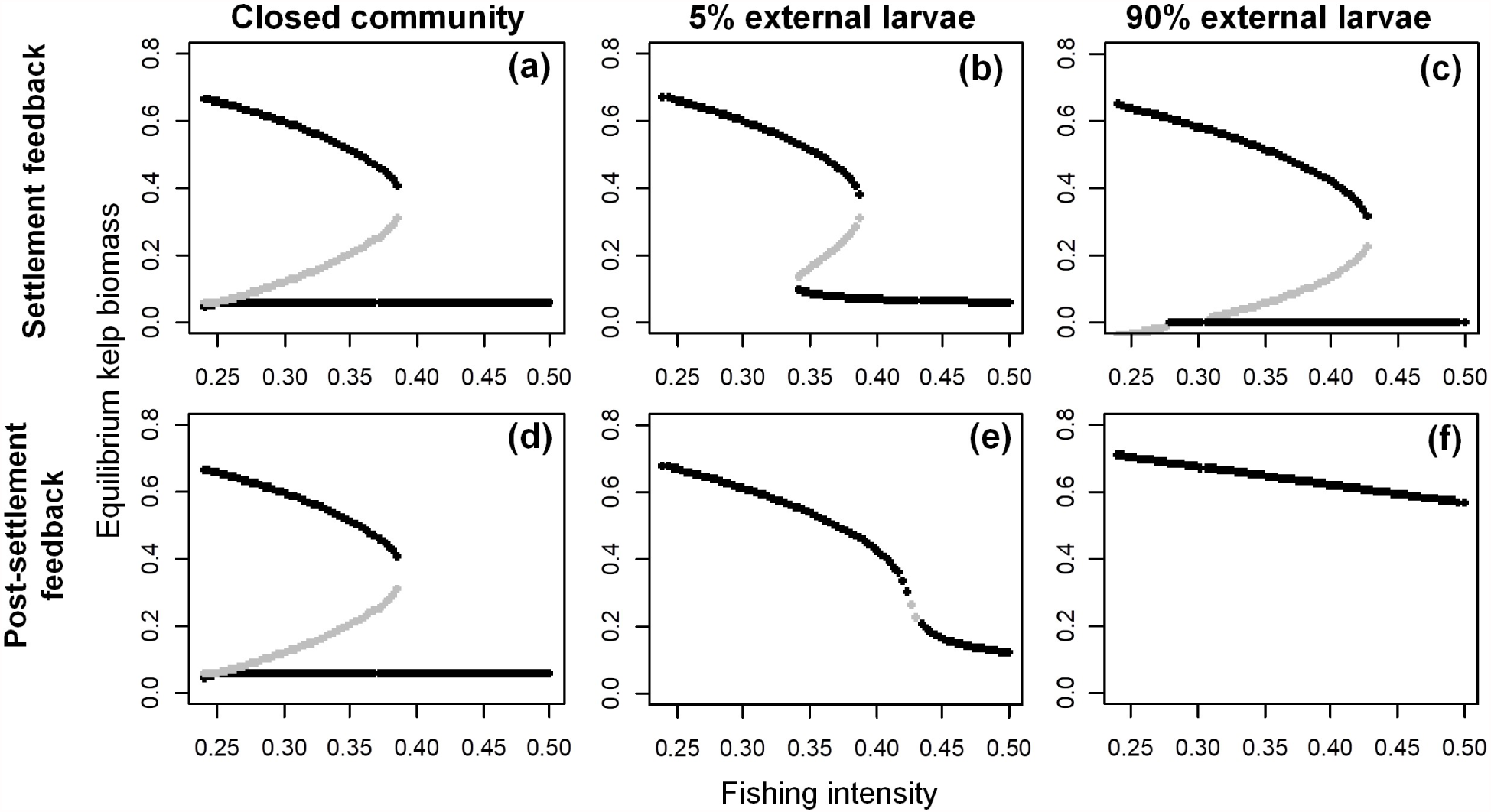
Effects of external larval supply on the presence of alternative stable states in deterministic community models of settlement (a-c) versus post-settlement (d-f) feedback loops. Closed (a, d), low (b, e) and high (c, f) external larval supply in each model correspond to *γ*_*U*_ = *γ*_*P*_ = 0, *γ*_*U*_ = *γ*_*P*_ = 0.05 and *γ*_*U*_ = *γ*_*P*_ = 0.9. Black lines represent stable equilibria and gray lines denote unstable equilibria. Kelp biomass in all cases (y-axes) is scaled to kelp carrying capacity. All parameters are as in Table 1.

## S3 Comparison to observed community scales

We use data from sites where kelp forests have been observed and where ample giant kelp habitat was available throughout the monitoring periods. In each year and at each site, data were collected in 15-50 1-5m^2^ samples (spaced over 0.002-0.5km^2^ area of reef) just after the peak growing season (June-August) when plants reach peak size and biomass (Cavanaugh et al. 2011). As in model simulations, observations exhibit two distinct kelp- and urchin-dominated states (Fig. S6a), but differ in that even low urchin densities can maintain low kelp densities because urchin grazing rates increase in barrens (Filbee-Dexter and Scheibling 2014). Nevertheless, in both our model and at monitoring sites urchin grazing creates barren community states with very low kelp densities. For consistency across surveys, we considered adults as > 1m tall individuals with ≥ 5 stipes (PISCO) or with haptera at or above the primary dichotomy (NPS); these classifications yielded analogous kelp densities where both surveys overlapped. We then classified each site in every year as forested or barren based on whether mature plant densities were above or below 1 ind. 100m^−2^; different choices of this density threshold yield qualitatively similar estimates of community scales. Importantly, in measuring spatial autocorrelation, we omit comparison of community states among (i) the same site in different years and (ii) among sites on Northern and Southern shores of the Channel Islands, which in a given year experience very different ocean conditions (analogously to estimating scale of urchin larval survival stochasticity).

Distributions of forest durations in the model and data remain similar when we compare only fully observed states for which initial and final years are known (D=0.14, *P* = 0.27). However, in this analysis distributions of barren durations were also similar among the model and data because this approach excludes lengthy, partially observed barren states (D=0.15, *P* = 0.24).

**Figure S6:**
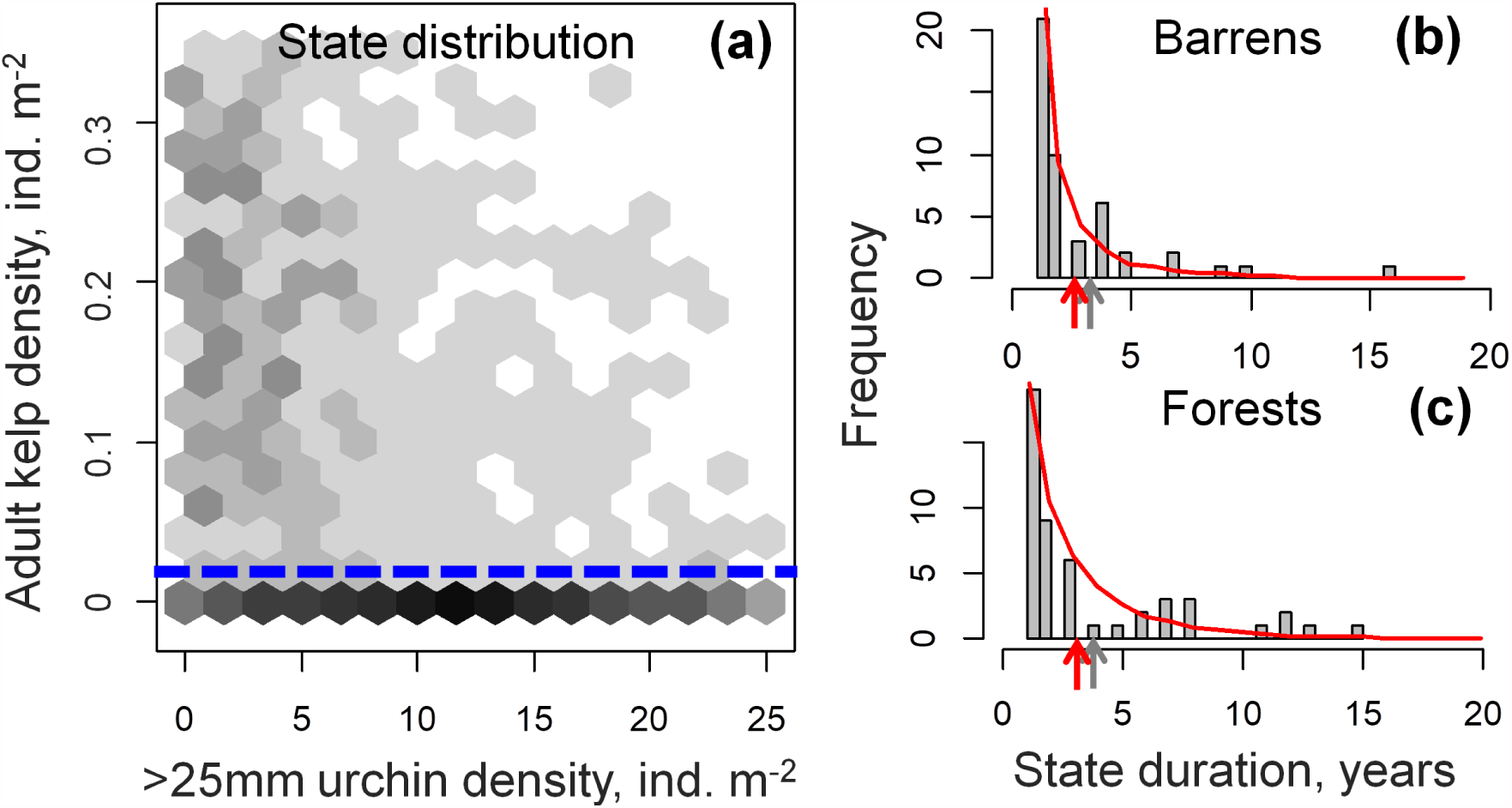
Distributions of observed community states across sites and years (a). Each bin represents mean densities of kelp (*M. pyrifera*) and urchins (> 25mm test diameter *S. purpuratus* and *Mesocentrotus franciscanus*) densities at each survey site and year (1336 total observations), and the dotted divides forest and barren states. Durations of fully observed barren (b) and forest (c) states in data (bars, frequencies) and model simulations (red).

## S4 Independence of initial conditions in stochastic models

To check that our analysis of spatially explicit models excludes the effects of lengthy transient dynamics, we compare community dynamics among model runs starting from spatially homogeneous initial conditions in a kelp forest versus a near-urchin barren state. Under the default parameterization, spatial models exhibit the longest transient dynamics, which disappear after the first 1000 years of model dynamics.

**Figure S7:**
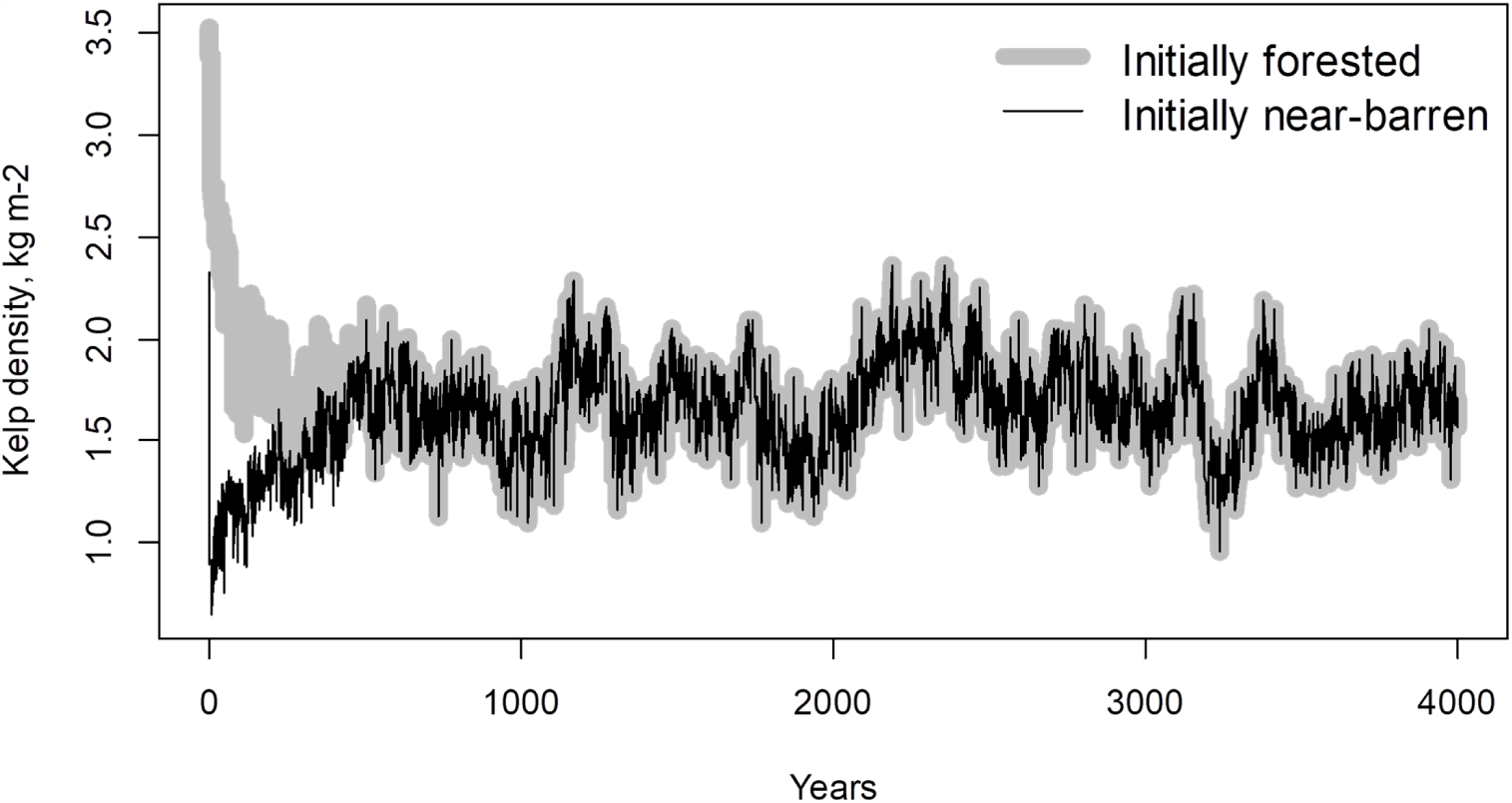
Effects of initial conditions (gray for initially forested and black for initially near-barren) on the temporal dynamics in kelp biomass aggregated across each system. After *t* ≈ 1000, models simulated under identical sequences of environmental stochasticity exhibit identical local dynamics. All parameters are as in Table 1.

## S5 Drivers of state shifts in stochastic models

Here we resolve the relative influence of kelp mortality and urchin larval survival variation in producing shifts among forested and barren states within local communities. Note that many shifts occur among ephemeral states (e.g., 1-2 year persistence) occur when communities near the threshold between the two states experience relatively minor disturbances. To resolve disturbances that have the strongest, lasting effects on community states, we instead focus only on community shifts that result in more persistent, ecologically relevant states (*>* 4 years). Overall, shifts among persistent states have a nearly three-fold greater Pearson correlation with urchin larval survival stochasticity (−0.132) than with kelp survival (0.046). Thus, barren formation typically occurred under favorable urchin larval survival stochasticity (*σ*_*U*_ (*τ*_*K*_) *>* 1; Fig. S8b) while forest recovery primarily corresponded with poor urchin larval survival (*σ*_*U*_ (*τ*_*K*_) *<* 1).

**Figure S8:**
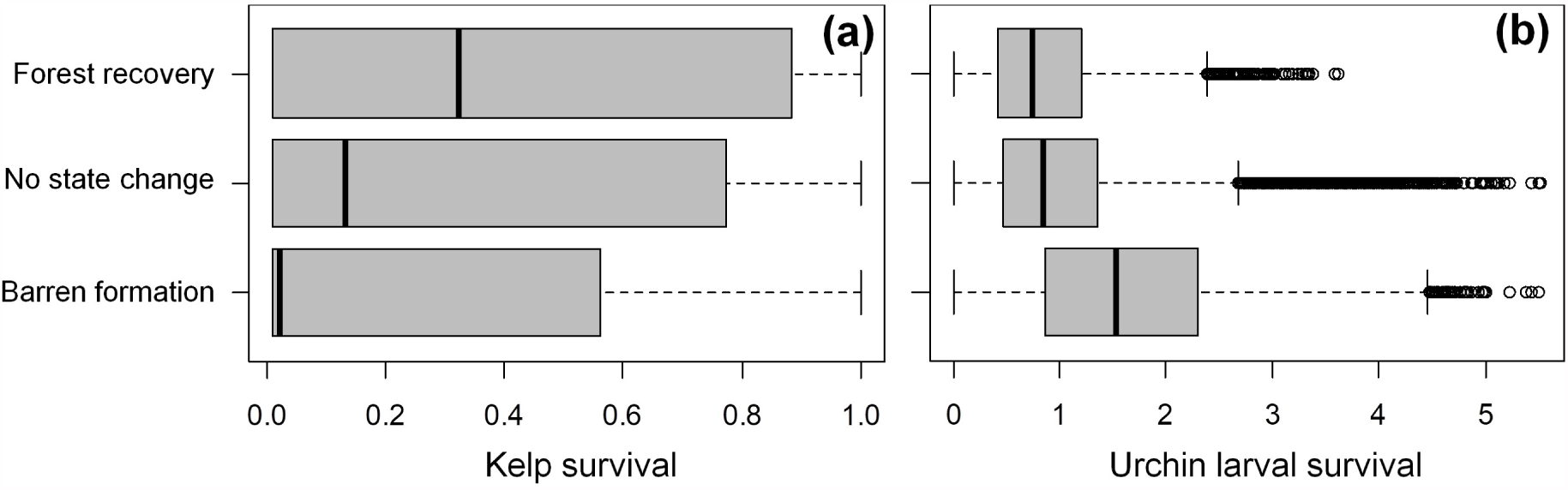
Effect of local kelp survival (a) and urchin larval survival (b) on the direction of change in persistent (*>* 4 years) local community states. All parameters are as in Table 1.

## S6 Sensitivity to recruitment facilitation

Repeating our analyses in Figures 3b and 4d, we find that alternative stable states occur at local rather than system-wide scales over a range of recruitment facilitation levels in our stochastic, spatial model. Overall, weaker facilitation feedbacks lead to a greater range of fishing intensities with intermediate frequencies of forest states that arise from the presence of localized alternative stable states. Note, however, that weaker facilitation reduces duration of both states due to a greater role of stochasticity, and the underlying deterministic model requires *f*_*c*_ > 0.5 for alternative stable stable states to be present (results not shown).

**Figure S9:**
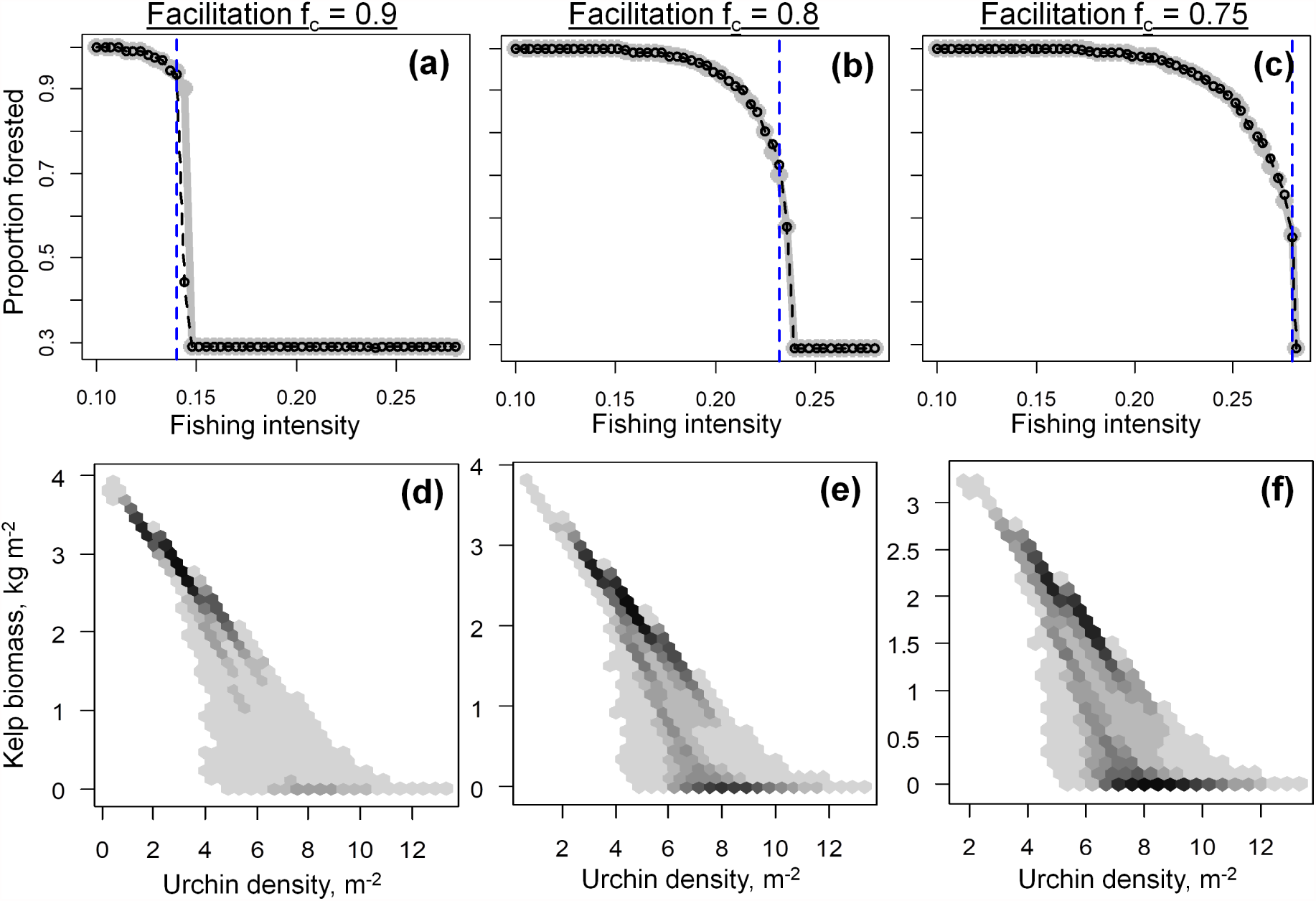
Long-term fraction of local communities that occur in kelp forest states for simulations starting from a forested (black, dashed lines) or near-barren (gray, solid lines) state (a-c). In every case, intermediate levels of forest cover that occur over a range of fishing intensities (blue dashed lines in a-c) correspond to bimodal distributions of local community states (d-f). All parameters except *f*_*c*_ are as in Table 1.

